# Model-guided gene circuit design for engineering genetically stable cell populations in diverse applications

**DOI:** 10.1101/2024.09.01.610672

**Authors:** Kirill Sechkar, Harrison Steel

## Abstract

Maintaining engineered cell populations’ genetic stability is a key challenge in synthetic biology. Synthetic genetic constructs compete with a host cell’s native genes for expression resources, burdening the cell and impairing its growth. This creates a selective pressure favouring mutations which alleviate this growth defect by removing synthetic gene expression. Non-functional mutants thus spread in cell populations, eventually making them lose engineered functions. Past work has attempted to limit mutation spread by coupling synthetic gene expression to survival. However, these approaches are highly context-dependent and must be tailor-made for each particular synthetic gene circuit to be retained.

In contrast, we develop and analyse a biomolecular controller which depresses mutant cell growth independently of the mutated synthetic gene’s identity. Modelling shows how our design can be deployed alongside various synthetic circuits without any re-engineering of its genetic components, outperforming extant gene-specific mutation spread mitigation strategies. Our controller’s performance is evaluated using a novel simulation approach which leverages resource-aware cell modelling to directly link a circuit’s design parameters to its population-level behaviour.

Our design’s adaptability promises to mitigate mutation spread in an expanded range of applications, whilst our analyses provide a blueprint for using resource-aware cell models in circuit design.

## 1 Introduction

Synthetic biology, which entails engineering living systems with useful functionalities by introducing new synthetic genes to cells, promises to tackle global challenges in medicine, industry and sustainability. However, synthetic biology is currently held back by several inherent challenges, such as non-modularity and evolvability of living systems, which make robust, predictable and durable performances difficult to achieve [1, 2].

A major factor contributing to these issues is the finiteness of the pool of cellular resources (most importantly for bacteria, ribosomes) which are shared between synthetic genes introduced by engineers and the host cell’s own native genes. On one hand, this complicates the design of so-called ‘circuits’ of synthetic genes that regulate each other to perceive, process and react to stimuli. Indeed, unlike electrical components, all gene circuit elements interact indirectly via the shared resource pool, which violates the engineering principle of modularity and makes it hard to predict a circuit’s performance based on the behaviours of its individual components observed in isolation [2, 3]. Moreover, high synthetic gene expression may significantly deplete the cellular resource pools, interfering with native gene expression and thus the cell’s growth and functioning. This imposes a burden on the host, which may have multiple downstream effects. For example, the changed cellular context (e.g. cell-wide variations arising from growth rate changes) can impact the circuit’s dynamics, making it even harder to predict or design [4, 5]. On the other hand, in an engineered cell population, mutants with defunct circuitry may experience a lesser burden and divide faster, leading to ‘mutation spread’ – that is, growth of non-functional sub-populations that eventually outcompete and displace original cells. This evolutionary pressure to get rid of engineered functionalitities significantly impairs the productivity and durability of biotechnologies [1].

To address these resource finiteness challenges, synthetic biologists have developed mathematical gene expression models that incorporate resource competition dynamics [2], as well as the host cell’s growth regulation mechanisms and their interplay with synthetic gene expression [4]. This has enabled more reliable predictions of the genetic circuit performance despite the non-modularity and context-dependence of biological systems. Furthermore, the insights provided by resource-aware model simulations and analytical derivations have facilitated the development of circuits rendering synthetic biology designs more modular and robust to resource competition [5].

To tackle resource competition’s population-level implications, several countermeasures to mutation spread have been proposed. Most straightforwardly, genes’ mutation probability can be decreased by codon optimisation or by removing the host cell’s homologous recombination machinery and transposable genetic elements which may disrupt DNA sequences, although this does not eliminate the possibility of mutations completely [6]. Alternatively, synthetic gene expression burden can be reduced to make the engineered cell’s growth defect less of a competitive disadvantage. This can be done either permanently by giving circuit genes weaker promoters and ribosome-binding sequences (though at the cost of often-desired high expression levels), or only when synthetic protein synthesis excessively stresses the cell, which can be detected by stress-response promoters used as negative feedback regulators [3]. In another strategy, engineered cells remain in a non-producing state (which little associated burden), and a fraction of the population in every generation is differentiated into a state with high synthetic gene expression [7]. Finally, to make a cell’s survival dependent on the presence of synthetic circuitry, a gene essential to cell growth is co-expressed with the synthetic genes of interest – potentially with these genes’ DNA sequences overlapping for even stronger coupling between their functioning. Loss of the useful synthetic genes’ expression to mutation would thus be accompanied by essential gene loss, making mutants unable to grow and outcompete engineered cells [6].

However, these extant mutation spread mitigation strategies can often have a restricted scope of applications and be cumbersome to adapt to new scenarios. Many of the aforementioned methods limit the maximum achievable productivity of biotechnologies, either because of keeping synthetic gene expression low by design (to reduce the burden and selective pressure against engineered cells) or due to confining the expression of genes of interest to just the subset of differentiated cells within the population. These bounds on a population’s synthetic capacity may be undesirable in applications such as biomanufacturing, where high product yields are required [3, 6]. Furthermore, most methods must be tailored for each application. The sequence-specificity of codon optimisation and essential gene overlapping mean that a given circuit’s DNA implementation has to be redesigned with these strategies in mind. Likewise, the addition of a differentiation switch and co-expressed essential genes requires case-specific additional circuitry. Having to design and synthesise new DNA sequences for every new circuit of interest is very time- and cost-intensive [8], which has limited the application of mutation spread mitigation methods in synthetic biology applications.

In this study we propose ‘the Punisher’, a novel circuit for countering mutation spread. Our versatile design deprives mutant cells of a growth advantage, yet can be easily adapted for different applications without re-engineering its DNA sequences. This work extends the applications of resource-aware gene expression modelling techniques [5], previously used for mitigating the effects of resource competition on a single-cell level, to countering undesired population-wide burden phenomena. A coarse-grained cell model provides a holistic view of resource competition between synthetic and native genes, as well as its effects on cell growth. Based on this single-cell modelling, we define a novel engineered cell population model that establishes a direct link between the circuit’s parameters and its population-level performance. Our analysis suggests scenarios in which the Punisher has advantages over the extant method of countering mutation spread by co-expressing synthetic and essential genes. Moreover, our modelling derivations inform the choice of the Punisher’s design parameters including those that can be adjusted without any genetic modifications in order to reuse the Punisher in novel applications.

## 2 Circuit design and analysis

### 2.1 Circuit description

Mutation spread occurs because cells in which synthetic genes mutate to become non-functional or non-expressed are rewarded by burden alleviation and the resultant growth advantage, which selects for such mutants and lets them eventually take over the population [1, 6]. To counter this, our circuit, shown in Figure 1A, replaces reward with punishment by detecting burden-alleviating mutations in its host *E. coli* cell and actively reducing cell growth in response. Responding directly to burden, regardless of which particular synthetic genes contribute to it, allows the Punisher to be easily reused across different applications.

**Figure 1:**
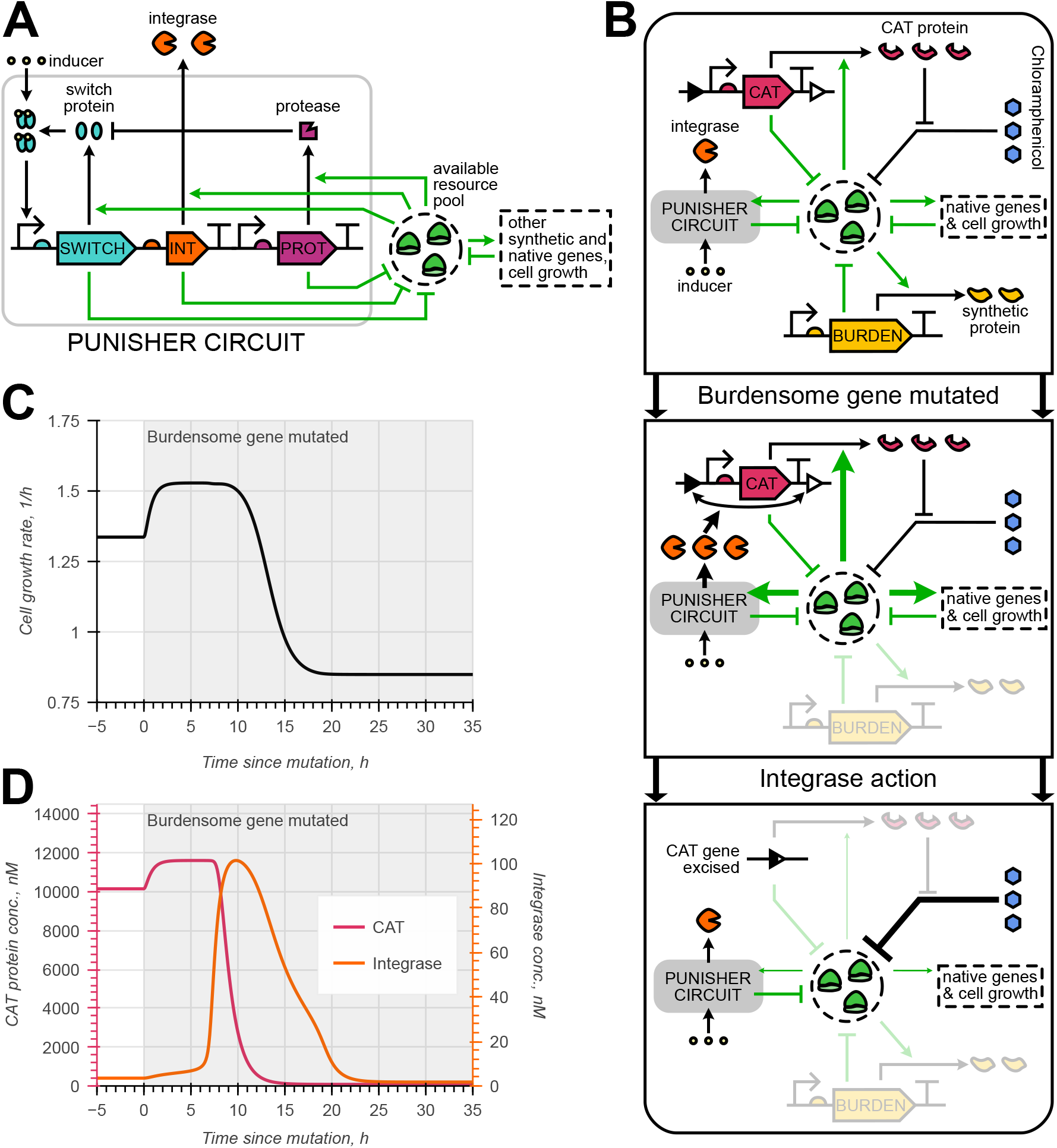
Basics of the Punisher’s functioning. **A** The Punisher consists of a self-activating switch gene and an integrase gene co-expressed with it. The switch protein is degraded by a synthetic protease and needs to be bound by a chemical inducer molecule in order to act as a transcription factor. All three proteins’ expression rates and steady-state concentrations depend on the availability of gene expression resources (ribosomes) in the cell. **B** Steps of the Punisher’s response to synthetic gene expression loss (arrow boldness reflects the strength of interactions). When the mutation of a burdensome synthetic gene frees up resources, increasing the mutant cell’s growth rate, more integrase is produced by the Punisher. Consequently, the antibiotic resistance gene CAT is excised by the integrase, so unhindered ribosome inactivation by the antibiotic chloramphenicol makes the available resource pool size decrease together with the mutant cell’s growth rate. **C–D** Simulation of the Punisher’s response to the mutation of a single constitutive burdensome gene expressed in the same host cell with it.

The Punisher’s detection component comprises the self-activating ‘switch’ gene, which encodes a transcription factor protein that, when bound by a chemical inducer molecule, promotes its own expression. Experimental studies show that such a protein’s concentration can converge to either a low- or a high-expression equilibrium depending on the burden experienced by the host cell [9, 10]. When all synthetic genes in the cell are functional, competition for gene expression resources is high, so the switch protein’s concentration *p*_*s*_ is prevented from reaching a high-expression equilibirum. Upon synthetic gene expression loss, more resources become available for the switch protein’s synthesis, so *p*_*s*_ starts converging to the high-expression fixed point and increases severalfold. Since the timescale of a biomolecular system’s dynamics is primarily determined by the rate of its removal from the system [11], we speed up the Punisher’s response by having the switch protein be not only diluted due to cell division, but also degraded by a synthetic protease [12, 13].

The punishment component is powered by an integrase protein co-expressed from the same operon with the switch gene. Once the integrase reaches a sufficiently high concentration, it excises the DNA sequence situated bewteen its cognate sites, which in our design flank a gene essential for cell growth. Hence, upon the detection of a mutation-induced rise in the switch’s and the integrase’s expression level, the essential gene is rapidly excised from its plasmid or genomic DNA [7, 14], which impairs the host cell’s growth. Importantly, the integrase’s action is irreversible, so a ‘punished’ mutant cell will remain slow-growing even if essential gene loss reduces resource availability, bringing the switch protein’s abundance back to its pre-detection level.

To illustrate our circuit’s operating principle, we consider the case of a single burdensome synthetic gene being found in the cell alongside the Punisher, depicted in Figure 1B. Here, the essential gene excised by the integrase is chloramphenicol acetyltransferase (CAT), an enzyme which degrades the ribosome-inactivating antibiotic chloramphenicol [15]. If chloramphenicol is present in the culture medium, CAT gene loss impairs translation in the cell and thus cell growth. Initially, only a few integrase molecules are present in the cell, so CAT’s concentration is high and steady. Upon the burdensome gene’s mutation, the cell growth rate’s rise is followed by a sharp increase in the integrase’s abundance. Consequently, the CAT gene is cut out and the cell growth rate slows down dramatically. Due to the excision’s irreversibility, the subsequent fall in the integrase’s concentration due to decreased ribosome availability does not recover the cell’s growth rate.

### 2.2 Model of circuit in the host cell context

Given that the Punisher reacts to changes in gene expression resource availability and curbs the host cell’s growth rate in response, informative modelling of the circuit must incorporate resource competition dynamics between native and synthetic genes, as well as capture the mechanisms determining the host cell’s growth rate. This is achieved using a coarse-grained resource-aware cell model [5] which, in addition to synthetic circuitry, explicitly considers the expression of a cell’s native genes and their interactions and regulation. Making sure that resource demand includes both synthetic and native genes contributing to it, our model can predict the cell’s growth rate and resource allocation, including when it is exposed to the ribosome-inhibiting antibiotic chloramphenicol, known to significantly influence the resource competition landscape [4]. Simultaneously, coarse-graining the cell’s native genes by their function into just several classes keeps the model simple, allowing the derivation of analytical relations describing the Punisher’s behaviour in Section 2.3.

Our ordinary differential equation (ODE) system is thus based on the resource-aware cell model from [5], modified to capture the intracellular chloramphenicol concentration and synthetic protease action as outlined in Supplementary Note S1.3. Host cell dynamics are described by Equations (1)– (6), and the behaviour of the synthetic gene set *X* is given by Equations (7)–(8).

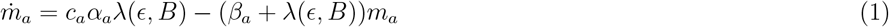

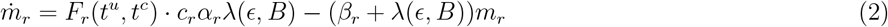

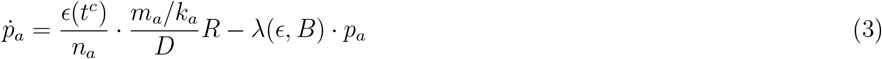

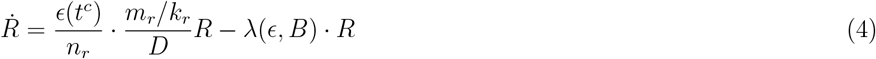

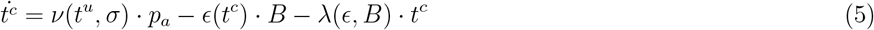

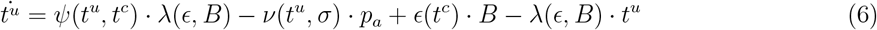

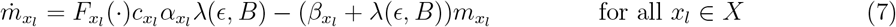

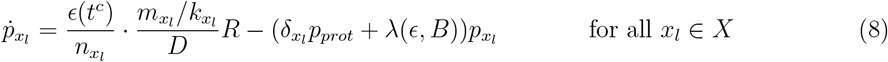

Two native genes classes are considered – ribosomal (*r*), responsible for protein synthesis, and metabolic (*a*), which produce aminoacyl-tRNAs *t*^*c*^ from uncharged tRNAs *t*^*u*^ using nutrients form the culture medium, whose quality is captured by the factor *σ*. The *r* and *a* gene classes are each treated as a single lumped gene whose mRNA concentrations is *m*_*r*_ or *m*_*a*_ respectively, and whose protein concentration is respectively *R* or *p*_*a*_. Out of *R nM* of ribosomes in the cell, *B nM* are actively translating. Both for native and synthetic genes, *c*_*j*_ is the gene *j*’s DNA concentration in the cell, *α*_*j*_ is its promoter strength, *n*_*j*_ is its length in amino acids, *β*_*j*_ is the mRNA degradation rate and *k*_*j*_ is the mRNA-ribosome dissociation constant, reflective of a gene’s ribosome-binding sequence (RBS) strength. Synthetic gene ODEs also include the (possibly zero) rate 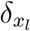 at which the Punisher’s protease (present in concentration *p*_*prot*_) degrades them, as well as the transcription regulation function 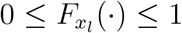 whose form and arguments depend on the particular gene considered. The functions *λ*(*ϵ, B*), *ϵ*(*t*^*c*^), *F*_*r*_(*t*^*u*^, *t*^*c*^), *ψ*(*t*^*u*^, *t*^*c*^) and *ν*(*t*^*u*^, *σ*) are given in Supplementary Table S2 and represent the cell growth rate, the translation elongation rate, the ribosomal genes’ transcription regulation function and tRNA synthesis and aminoacylation rates, respectively.

The variable *D* (specified in Equations (9)–(10)) is the ‘ribosomal competition denominator’ capturing the availability of translational resources in the cell. Besides the already-explained variables, it is affected by the mass fraction of housekeeping (i.e. not metabolic or ribosomal) native proteins in the cell 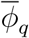, assumed constant [5, 15], and the concentration *h* of chloramphenicol in the cell, which binds and inactivates ribosomes with a dissociation constant *K*_*D*_.

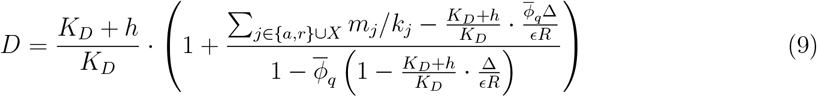

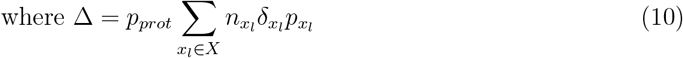

The synthetic gene set *X* includes the synthetic circuitry whose mutation the Punisher aims to penalise – explicit ODEs characterising different setups that we consider are provided in Supplementary Note S1.5. Moreover, *X* includes the Punisher’s switch, integrase, protease and CAT genes (*X* ⊇ *{s, i, prot cat}*) described by Equations (11)–(19), where *I* is the share of switch proteins bound by the chemical inducer molecules, *η*_*s*_ and *K*_*s*_ are the cooperativity and dissociation constant for the switch protein’s binding to the switch gene’s DNA, and 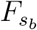 is the switch gene promoter’s baseline activity without transcriptional activation.

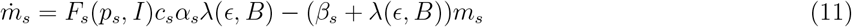

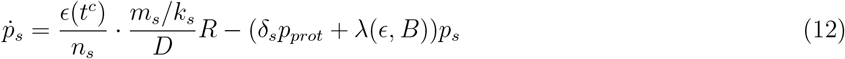

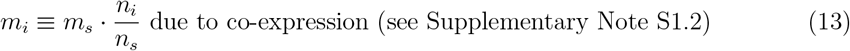

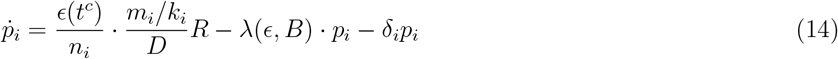

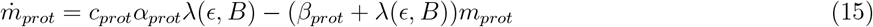

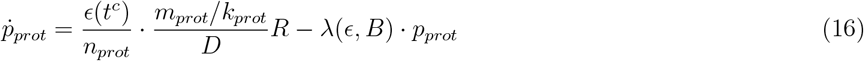

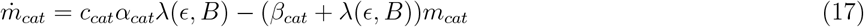

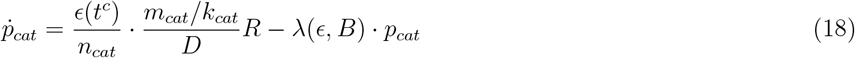

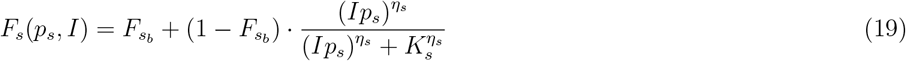

While all other genes’ concentrations remain constant, CAT gene DNA can be excised by the integrase in a reversible strand exchange reaction followed by an irreversible conformation change. We model these reactions using Equations (20)–(21) derived in Supplementary Note S1.4 based on an experimentally parameterised serine integrase action model from [14] and [16]. Here, *c*_*cat*_ and *c*_*LRi*_ are the concentrations of the CAT gene DNA before and after strand exchange (the former being the functional gene copy number), *K*_*bI*_ is the integrase-DNA dissociation constant, 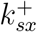 and 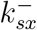 are the forward and backward strand exchange rates, and *k*^*conf*^ is the conformation change rate.

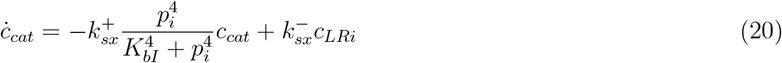

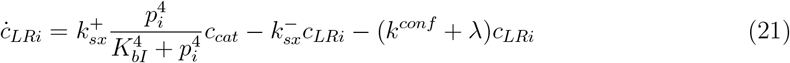

If the CAT protein’s rate of binding and degrading chloramphenicol is captured by the affinity constant *K*_*C*_, the antibiotic’s intracellular dynamics can be predicted by Equation (22), where *κ* being the rate of chloramphenicol’s diffusion through the cell membrane [15, 17].

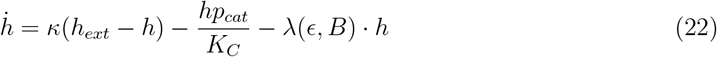

We use these ODEs to simulate the case of a single constitutive synthetic gene being present in the cell alongside the Punisher, described in Section 2.1, Figure 1B and Supplementary Note S1.5.1. The obtained trajectory in Figure 1C–D matches our expectations for the Punisher’s performance. The same behaviour is reproduced by stochastic cell model trajectories, simulated according to the hybrid tau-leaping method described in [5] (Supplementary Note S2.1). The code for these and all other simulations was implemented in Python 3.12 using the JAX 0.4.23 package to enable efficient parallelised computation [18], and is available at github.com/KSechkar/punisher.

### 2.3 Switching threshold identification and tuning

As described in Section 2.1, the Punisher detects synthetic gene mutations if they reduce the burden experienced by the host cell below the threshold value at which the switch gene transitions from a low-expression to a high-expression equilibrium. Thus, predicting whether the Punisher will function correctly requires understanding what determines its switching threshold and how to adjust it. Moreover, although multiple different synthetic circuits may burden the cell to a similar extent (making it possible to use the same implementation of the Punisher with either of them), synthetic gene expression burden may vary depending on the circuit’s architecture and the cell culture conditions. Therefore, for the Punisher to be deployable in a new scenario where synthetic gene mutations alleviate burden to a different extent, guidance on how to adjust the Punisher’s switching threshold is also required. Such design insights are enabled by our coarse-grained cell model’s simplicity and amenability to analytical derivations.

As we show in Supplementary Note S3, with a few realistic simplifying assumptions our coarsegrained resource-aware cell model allows to quantify the steady-state burden of expressing gene *j*, be it native or synthetic, as

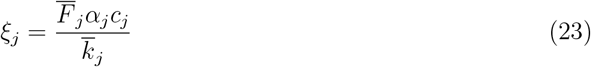

where 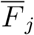 is the steady-state value of its transcription regulation function. The total gene expression burden sensed by the Punisher is the sum of these burden factors over all genes except for the burden imposed by the Punisher’s switch and integrase genes themselves:

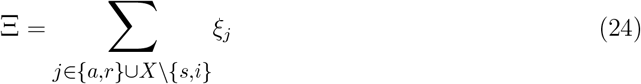

The cell model can then be solved at steady state to find the value of *F*_*s*_ required to achieve a given steady-state switch protein concentration *p*_*s*_ for a given burden. This yields the 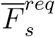 value (Equations (25)–(26)), with 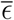 and 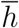 standing for the steady-state translation elongation rate and intracellular chloramphenicol concentration, *M* being the host cell’s total protein mass, and 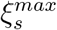 and 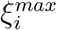 representing the maximum (i.e. calculated for *F*_*s*_ = *F*_*i*_ = 1) burden of expressing the switch and the integrase proteins.

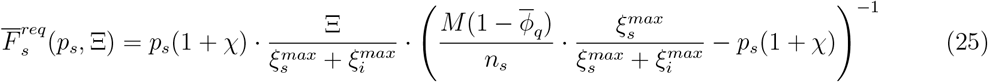

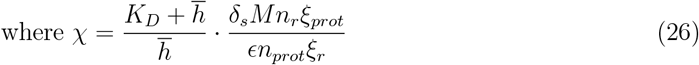

However, the actual value of the switch gene’s transcription regulation function is given by a Hill function in Equation (19). The switch protein concentration is therefore in steady state if and only if

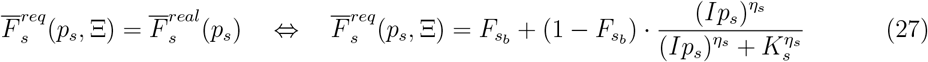

i.e. the real transcription regulation function’s value is equal to the value required to achieve equilibrium. Graphically, this can be understood as the blue and the black curves intersecting in Figure 2A. The plot also reveals the fixed points’ stability: the switch protein’s concentration decreases when the actual transcription rate is below that required to keep *p*_*s*_ steady, and increases in the opposite case.

**Figure 2:**
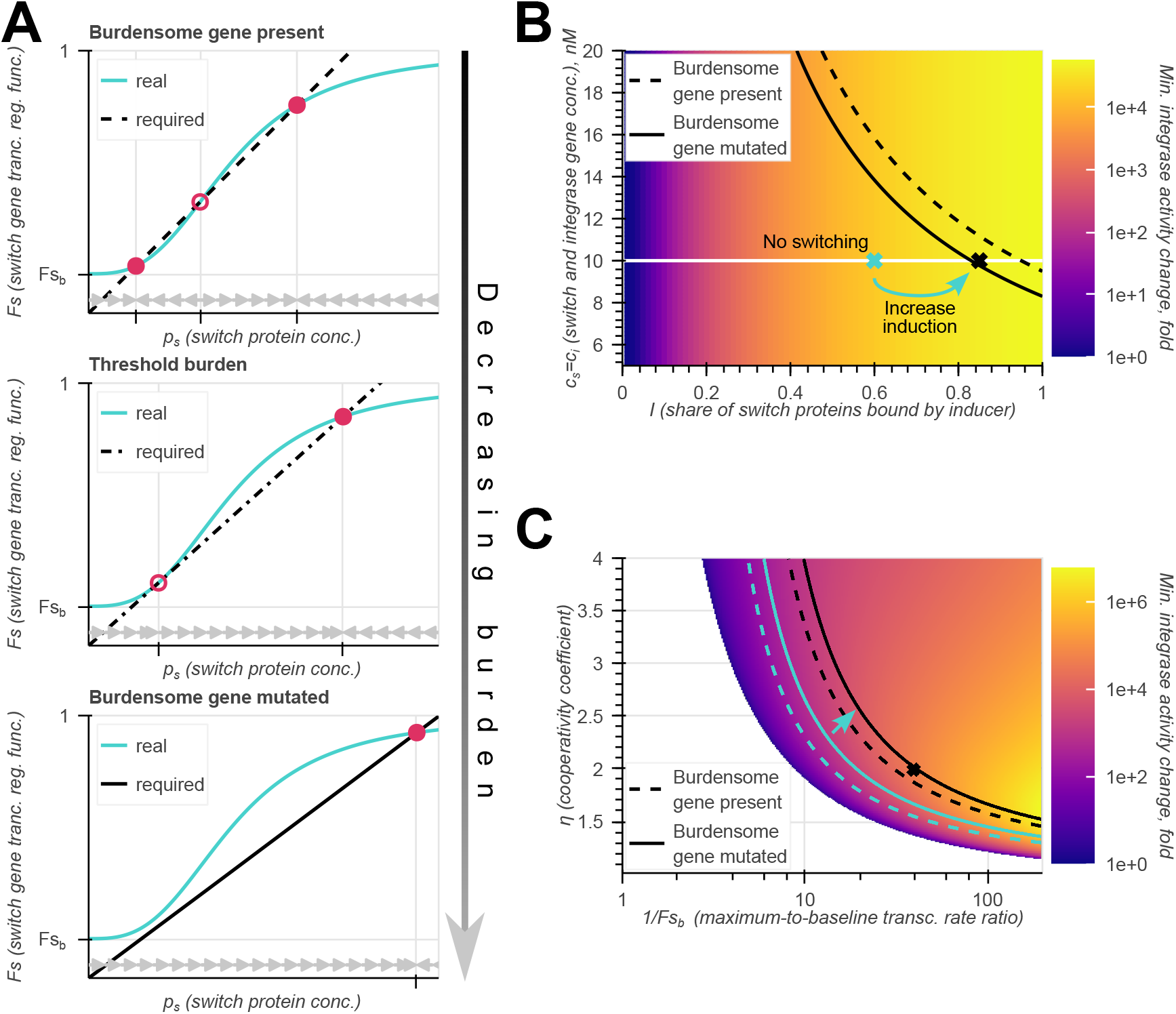
Emergence, identification and tuning of the Punisher’s switching threshold. **A** Required 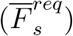 and real 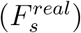 values of the switch gene’s transcription regulation function calculated from the switch protein concentration *p*_*s*_ for the case of all synthetic genes being functional (top), at the threshold burden 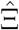 (middle), and when synthetic gene expression has been lost (bottom). Circles mark the system’s fixed points – filled for stable, empty for unstable. Grey arrows mark the direction of convergence of *p*_*s*_. **B** Heatmap of the minimum fold-change in the integrase’s DNA-cutting activity as a function of the switch and integrase genes’ DNA concentration *c*_*s*_ = *c*_*i*_ and the share of switch protein bound by the inducer *I*. Here and in (C), the dashed and solid black lines respectively represent parameter combinations for which the switching threshold 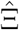 is equal to burden with and without the expression of the single constitutive synthetic gene. The Punisher will function correctly for all parameter combinations between these two lines, which define the acceptable parameter region. The black cross represents the parameter combination used in simulation in Figure 1. If initially the Punisher’s parameters do not make it switch on upon synthetic gene expression loss (e.g. blue cross at *I* = 0.6), the inducer’s concentration in the medium can be tuned to achieve the desired behaviour (blue arrow). **C** Heatmap of the minimum fold-change in the integrase’s activity as a function of the switch gene’s cooperativity coefficient *η*_*s*_ and the maximum-to-baseline promoter activity ratio 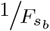. In the blank region, switching does not occur at any value of Ξ. Likewise to (B), the black lines denote the parameter combinations for which 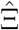 is equal to burden with and without synthetic burdensome gene expression, delineating the acceptable parameter region. Meanwhile, the blue lines mark the acceptable parameter region for *I* = 0.6, which can be shifted by increasing *I* (blue arrow) so as to cover the 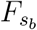 and *η* value combination used in Figure 1 (black cross).

Equation (25) indicates that Ξ defines the black line’s gradient in Figure 2A. Hence, for large gene expression burden this line rises steeply and crosses 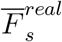 at a low *p*_*s*_ value, producing a stable equilibrium (Figure 2A, top). As Ξ is reduced, the curve’s slope becomes gentler, until it touches 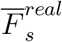 from below, producing a saddle-node birfurcation with a saddle node at low *p*_*s*_ and a stable fixed point to its right (Figure 2A, middle) [10]. If burden is further decreased below this threshold, a single high-expression equilibirum remains (Figure 2A, bottom). Crucially, since the switch gene and the integrase are co-regulated, the low-expression equilbirum stands for low integrase abundance and CAT gene excision rate (*p*_*i*_ ≪ *K*_*bI*_), whereas the high-expression fixed point corresponds to high integrase concentration and thus high essential gene excision rate.

Therefore, the Punisher penalises synthetic gene mutation when it brings the burden Ξ below the bifurcation threshold 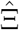. Retrieving this threshold value from the Punisher’s design parameters (see Suppplementary Note S3.4) and comparing it to burden before and after synthetic gene mutation allows to predict whether the Punisher will become activated in a mutant cell. Moreover, a lower bound on the change in integrase activity upon the Punisher’s triggering can be found according to Supplementary Note S3.5. This reveals how the Punisher’s design parameters define its performance. For instance, Figure 2B shows its dependence on the switch and integrase genes’ copy number and the inducer’s concentration in the culture medium. Meanwhile, Figure 2C demostrates the effect on the cooperativity coefficient *η*_*s*_ and the baseline promoter activity 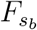, which are key determinants of a self-activating gene’s equilibria [19].

The dependence of 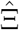 on *I* is of particular significance. While all other parameters are determined by the synthetic DNA sequence during construct design, the inducer’s concentration can be easily tuned by adjusting the chemical inducer’s concentration in the culture medium. Graphically, it stands for moving along the white line in Figure 2B to position 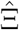 between the expected pre- and post-mutation burden levels. In plots for other parameters, such as Figure 2C, changing the chemical induction is equivalent to moving the acceptable parameter region to make it cover a given parameter combination. Therefore, without any re-engineering of its genetic components, the same DNA implementation of the Punisher can be reused in different conditions and with sundry synthetic gene circuits of varying burdensomeness simply by adding different amounts of the inducer to the medium.

## 3 Example application

### 3.1 Performance simulation

We now simulate the Punisher’s deployment alongside more complex synthetic gene circuitry than a single constitutive gene in Figure 1. We consider the cell hosting two synthetic toggle switches as shown Figure 3A, described with ODEs in Supplementary Note S1.5.2. A toggle switch comprises two genes that repress each other’s expression; this repression’s strength can be modulated by adding chemical inducers to the medium. Therefore, a toggle circuit exhibits bistability, since either of the toggle’s two genes can be highly expressed whilst repressing the other gene’s expression. Pulses of inducer concentration ‘flip’ the toggle from one equilibrium state to the other, in which it stays until the next flipping [20]. Several toggle switches may be required in the same cell if it needs to simultaneously ‘remember’ several different inducer pulses occurring in the past.

**Figure 3:**
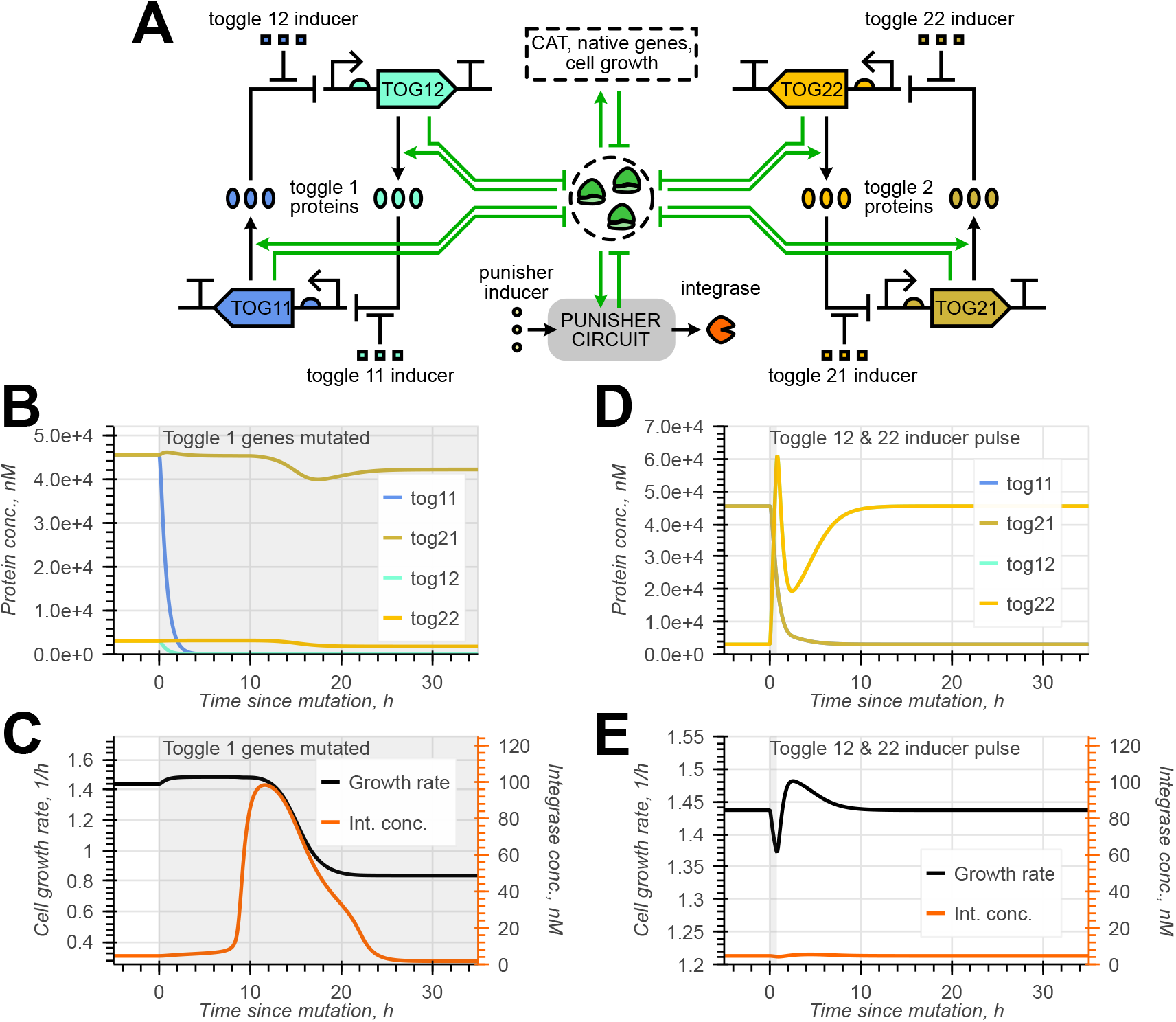
Using the Punisher with two synthetic toggle switch circuits. **A** Schematic of the two toggle switches being expressed in the same cell as the Punisher circuit. **B–C** Simulation of the Punisher’s response to loss of expression of the first toggle’s both genes. **D–E** Simulation of the Punisher’s response to the flipping of one toggle by a transient inducer concentration pulse.

Importantly, loss of the expression of a single synthetic gene in a network does not always alleviate burden [1]. Here, too, mutating one gene in a toggle switch means that the other gene is no longer repressed and is thus expressed even more actively, increasing burden and slowing down cell growth (Supplementary Note S2.2). Therefore, single-gene mutants have a growth disadvantage compared to the original engineered cells and present no risk of outcompeting them in the population. Losing both genes of a toggle switch, conversely, may significantly increase resource availability. In line with our derivations in Section 2.3, the inducer level should thus be changed to set *I* = 0.87 and position the Punisher’s switching threshold between the burden of expressing both toggle switches and that of expressing a single toggle switch. As we observe in Figure 3B-C, the Punisher then successfully detects the loss of a toggle and reduces cell growth in response.

When a synthetic circuit is out of steady state, the burden caused by it can vary over time. For instance, when a toggle switch is flipped, the overall expression of its genes may momentarily dip. Simultaneous flipping of both toggle switches can temporarily reduce burden to an extent close to that caused by mutating one toggle switch. Nonetheless, the Punisher can reject (i.e. not respond to) such transient disturbances (Figure 3D-E). Provided that the burden reduction lasts for a sufficiently short time period, the Punisher still may not leave the low-expression equilibirium’s basin of attraction by the time the burden returns to its original high value; therefore, the Punisher goes back to the off-state without triggering essential gene excision. Knowing the expected duration of disturbance to be filtered out, the Punisher’s switching timescale can be adjusted accordingly, e.g. by shifting the bifurcation threshold: as shown in Supplementary Figure S6A–B, the closer it is to the post-mutation burden, the slower is the Punisher’s reaction. Alternatively, assuming that the switch protein’s degradation rate can be adjusted – either by changing its protease affinity or altering the protease’s expression level – the timescale of response to mutation can be tuned without shifting the detection threshold as shown in Supplemenatry Note S3.6 [21].

### 3.2 Comparison to alternative mutation spread mitigation strategies

In certain applications, especially when relatively complex circuits are to be retained by cell populations, the burden-sensing nature of the Punisher’s response makes it advantageous over the extant mutation spread mitigation strategies. An instance of this is the task of penalising the mutations of two toggle switches, fulfilled using the Punisher in Section 3.1. Here, we compare our design’s performance in this case to that of the co-expression method discussed in Section 1 and [6], where the synthetic gene to be retained by the cell population is expressed together with a gene essential for growth, so mutating the former also disables the latter.

We simulate the scenario depicted in Figure 4A and described with ODEs in Supplementary Note S1.5.3. Here, the cell hosts two synthetic toggles whose genes (for symmetry, all four of them) are co-transcribed in the same operon with the antibiotic resistance gene CAT. Despite being translated from the same mRNA, essential and toggle gene expression may not be identical due to differences in post-transcriptional behaviour, which we capture by considering a wide range of possible CAT gene RBS strengths [4, 22, 23]. As revealed by Figures 4B, for most synthetic gene mutation combinations the punisher slows down mutant cells’ growth more than or at least closely to the best-possible co-expression setup.

**Figure 4:**
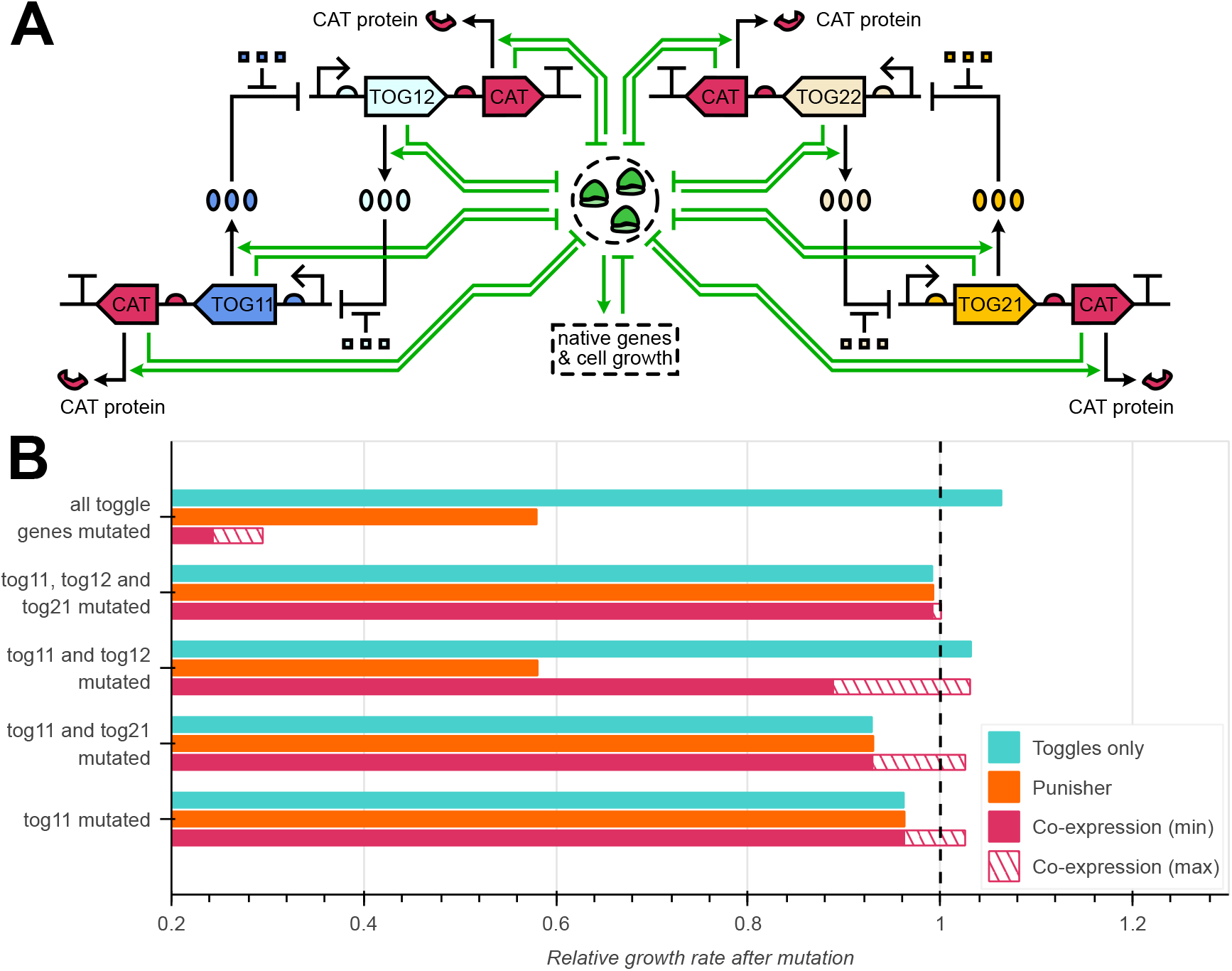
Comparing the Punisher’s and essential gene co-expression’s performance for penalising the mutations of two synthetic toggle switches. **A** Schematic of a pair of two toggle switches, all of whose genes are co-expressed with essential CAT gene copies. **B** The host cell’s steady-state growth rates (obtained by simulating the system for 50 *h*) with certain toggle genes mutated relative to its growth rate with all synthetic circuitry fully functional. The two toggle switches were assumed to be present in the cell without any mutation-penalising circuitry, alongside the Punisher as shown in Figure 3, or co-expressed with the CAT gene as per (A). For the latter, CAT mRNA-ribosome association rates 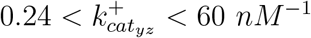*· h*^−1^ were considered [4, 22, 23], and the minimum and maximum relative growth rates in this RBS strength range are shown.

More significantly, in some parameter regimes CAT gene co-expression in fact promotes the growth of cells with undesirable mutations, which without they would be unable to take over the cell population. This is because the co-expression method operates with not the burden which slows down cell growth, but rather mutations themselves, whose influence on the growth rate may be less straightforward [1, 6]. For instance, as shown in Supplementary Note S2.3, mutating one gene in the toggle upregulates its counterpart, formerly repressed by it. This can increase synthetic protein expression levels above its original values, elevating the expression burden – which in this case is desirable as it selects against mutant cells. However, if this newly derepressed gene is co-expressed with an essential gene, the benefit to cell viability from increased essential protein levels can counteract this useful additional burden, potentially even producing mutant cells that grow faster than their unmutated progenitors. In contrast, the Punisher only reacts to synthetic gene expression loss when it does reduce burden and increase cell growth rates, removing the possibility of responses that actively (and undesirably) promote mutation spread.

Engineering essential gene co-expressions can also be more cumbersome than using the Punisher to counter mutation spread. First, a circuit may require multiple co-expressions (e.g. in our case, we use four of them), implementing which in DNA is time- and cost-intensive [8]. To apply the co-expression method to another circuit, this DNA engineering step would have to be repeated anew. Second, only certain parameter regions (e.g. RBS strengths 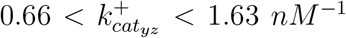.*h*^−1^ in our case) yield co-expression setups which do not risk inadvertently promoting mutation spread. Meanwhile, parameter tuning for this method can be challenging. Adjusting RBS strengths requires DNA editing, while the design space for ribosome-binding sequences may be restricted if the essential gene is not merely co-expressed, but actually overlaps with a synthetic gene’s sequence for stronger protection against mutations [6]. CAT genes’ transcription, the most commonly tuned expression step in synthetic biology, is governed by the toggles’ promoters and thus is not easily adjusted. Less trivial post-transcriptional regulation circuitry, which can be complicated to implement, may therefore be necessary to implement a functional essential gene co-expression setup.

Conversely, the same DNA implementation of the Punisher can be reused in different applications. Although our design, too, is only effective within a certain parameter region (see Figure 2), the switching threshold can be adjusted without costly genetic interventions by changing the concentration of its inducer in the medium as shown in Section 2.3.

## 4 Population simulations

### 4.1 Population model definition

Previous sections have demonstrated that the Punisher can detect synthetic gene mutation-induced changes in the burden experienced by a single cell, curbing its growth in response. However, the Punisher’s ultimate purpose is to slow down the outcompetition of engineered cells by mutants in a cell population, which calls for a population-level model that would allow to gauge our design’s performance *in silico* before testing it experimentally. To this end, we propose a model of a population of cells in a bioreactor, defined for the case of the Punisher being used to favour the retention of a single synthetic burdensome gene as described in Section 2.1.

Given the very high number of cells present in a typical bioreactor [1], an agent-based approach, capturing each individual cell’s behaviour with a separate model and modelling different cells’ interactions between each other [24], is computationally intractable. Instead, population-scale models, frequently used in evolutionary and synthetic biology [7], can classify cells as members of sub-populations according to their genetic state – that is, which synthetic genes remain unmutated and functional – and treating the abundance of each cell type as a variable modelled with ODEs. The combination of this approach with resource-aware cell modelling was pioneered by Ingram and Stan [1]. However, their model assumed identical internal state dynamics (captured by an ODE cell model) for every cell in a given genetic state. This is unsuitable for modelling genetic circuits with dynamics like that of the Punisher, which takes time to become activated upon synthetic gene mutation, hence the state of a recently mutated cell’s circuitry being different to that for a cell mutated long ago.

We thus further subdivide genetically identical cells according to the state of the circuit in each cell, and characterise the rates of switching between circuit states by simulating our resource-aware single-cell model (Figure 5A). The resultant model is depicted in Figure 5B and described and parameterised in Supplementary Note S3. The cell population is split into 2^4^ = 16 genetic states based on the functionality of: the burdensome gene to be retained (*B*); the switch and integrase genes (*S*, treated as one gene due to being co-expressed); the synthetic protease gene (*P*); and the CAT gene (*C*). If a given gene is mutated, the genetic state’s index has a ^*′*^ mark after the corresponding letter. Each genetic state can have three possible states of the Punisher – the high-expression equilibirum (*H*), the low-expression equilibrium (*L*) or else no integrase and switch proteins present (0), e.g. when the Punisher has mutated and all its previously synthesised proteins have been removed from the cell. If some equilibrium does not actually exist in a given genetic state (like the low-expression equilibrium when the burdensome gene is non-functional), the corresponding cell state’s properties are assumed identical to those of its closest unmutated progenitor according to Supplementary Table S7. We thus have 16 *×* 3 = 48 variables in total, each representing the abundance of cells with a given genetic state and a given state of the Punisher. The index *j* = *B*^*′*^*SPC* : *H*, for example, stands for the cells whose Punisher circuit is in a high-expression equilibrium, where only the synthetic burdensome gene *B* has been mutated.

**Figure 5:**
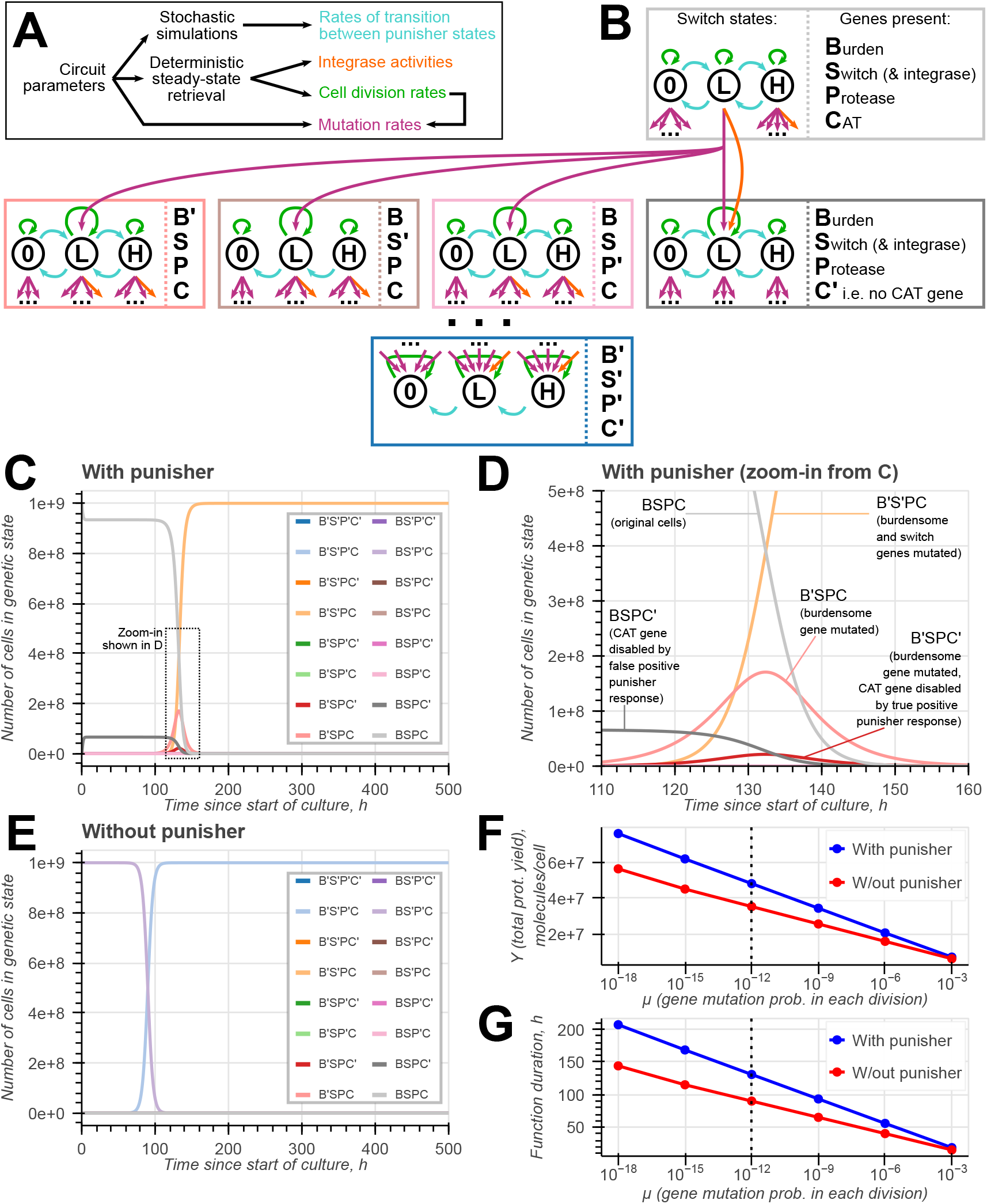
Definition and simulation of the model of a population of cells engineered with a synthetic burdensome gene and the Punisher circuit. **A–B** The population is split into subpopulations of cell based on their genetic state (i.e. which synthetic genes are still present or have been mutated) and state of the Punisher’s switch gene (zero, low or high expression). The rates of transition between the Punisher states are found by stochastic simulations of our single-cell model. The cell can change its genetic state either due to mutations during cell divisions or due to integrase action, the rates of which are found by deterministic single-cell simulations. Each subpopulation replenishes itself by cell division, whose rate is also found deterministically. **C–E** Simulations of the population model with and without the Punisher circuit. Every synthetic gene assumed to mutate with a probability of *µ* = 10^−12^ with every cell division. **F–G** Total protein yield per cell and function duration of engineered cell populations with and without the Punisher for different synthetic gene mutation probabilities per cell division. The dotted line represents *µ* = 10^−12^ in (C–E).

Each cell in a state *j* divides at a rate *λ*_*j*_ to produce two daughter cells, usually in the same state – however, with probability *µ* a still-functional gene may become mutated, making the daughter cells contribute to another genetic state’s total cell count. A genetic state transition can also be caused by the integrase excising the CAT gene at a rate dependent on the integrase’s concentration (i.e. on the Punisher’s state). Genetic changes do not directly affect the Punisher’s state; instead, the Punisher’s transition rates are fixed but dependent on the genetic state the cell is in. We determine them by simulating our resource-aware single-cell model stochastically [25]. This can capture not only ‘true positive’ detection of mutations predicted by ODE simulations, but also the ‘false positive’ activations of the circuit due to the stochasticity of gene expression, as well as the possible effects of stochasticity on the timescale of the self-activating switch gene’s expression dynamics [19].

In summary, the evolution of cell counts in the bioreactor is given by:

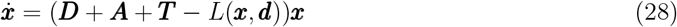

where ***x*** is the 48-dimensional vector of cell counts by the state and ***d*** is the vector of corresponding cell division rates. In the matrix ***D, D***_*j,l*_ is the rate of state *j* cell counts changing due to state *l* cells dividing – whilst the entry ***D***_*l,l*_ (i.e. cells dividing to produce cells of the same type) predominates, there may be other positive elements due to the possibility of gene mutations during cell division. In the matrix ***T***, ***T*** _*j,l*_ is the increase in state *j* cell counts as a result of the Punisher’s transitions from the state *l* (hence ***T*** _*l,l*_ being non-positive ∀*l*). Likewise, ***A***_*j,l*_ is the rate of genetic transitions from state *l* to state *j* due to integrase action. The definitions for ***d, D, A*** and ***T*** can be found in Supplementary Note S3.3. Finally, if we assume that the bioreactor is a turbidostat that keeps the overall cell abundance constant [15], it dilutes all cells at the same rate of

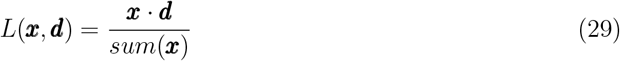

### 4.2 Population simulation results

To gauge the Punisher’s performance on a population level, we integrated Equation (28), starting at the initial condition where all cells in the bioreactor had all synthetic genes unmutated and the Punisher in a low-expression equilibrium, i.e.

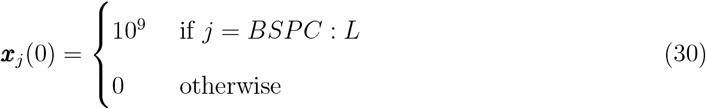

For comparison, we simulated a population of cells lacking the Punisher’s switch, integrase and protease (hence the zero switch and integrase protein level), but still hosting the same burdensome and CAT genes. This is equivalent to having the initial condition

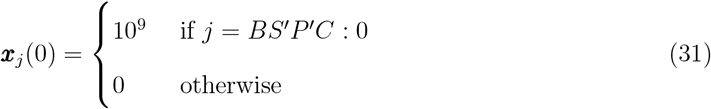

Representative simulated trajectories for the cell population dynamics with and without the Punisher, plotted in Figure 5D–E, qualitatively confirm that our design allows the cells with a functional burdensome gene to dominate the population for longer. A quantitative measure of an engineered cell population’s productivity is its rate of burdensome protein synthesis per cell Θ, defined in Equation (32), where *p*_*b*_(*j*) is the burdensome protein content of a cells in state *j*, found by simulating our ODE single-cell model [7].

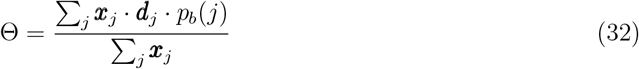

To gauge the engineered cells’ total productivity over time, Θ can be integrated over the culture duration *t*_*cult*_ = 500 *h* to find the total burdensome protein yield per cell *Y*. Moreover, Θ can be tracked over time to find a population’s function duration *τ*, which we define as the time for which *H* remains above 50% of its maximum value. Plotting these metrics in Figure 5F–G, we see that the Punisher increases the robustness to mutations of the cell population’s productivity roughly 1.5-fold over a wide range of gene mutation rates.

The Punisher’s observed beneficial effect on the engineered population’s function duration is explained by the cells having to accumulate two mutations, instead of just one, in order to gain a substantial growth advantage. As shown in Figure 5D, cells with just the burdensome gene mutated (state *B*^*′*^*SPC*) multiply very slowly, becoming noticeable only at *t* ≈ 100 *h* – this is because the Punisher detects the reduced burden in them and disables their CAT gene, with the resultant *B*^*′*^*SPC*^*′*^ genes dividing very slowly and being rapidly diluted out of the bioreactor. The rapid displacement of the original engineered cells therefore only becomes possible when a (rare) *B*^*′*^*SPC* mutant also mutates the Punisher’s switch and integrase genes to escape penalisation.

In practice, delaying of mutation spread’s onset with the Punisher may prove even more beneficial for the cell population’s productivity than predicted, which can be understood through the lens of the clonal interference phenomenon [26]. Besides synthetic gene mutations, cells in a bioreactor may undergo native gene mutations that confer them with a growth advantage. Therefore, the Punisher’s extension of the time throughout which original engineered cells predominate in the population increases the chance of such beneficial native gene mutations first arising in the cells with fully functional synthetic circuitry. This may allow them to outgrow undesirable cells with a mutated synthetic burdensome gene.

## 5 Discussion

In usmmary, we have leveraged known resource competition phenomena to design a novel versatile biomolecular controller that counters mutation spread in engineered cell populations. Numerical simulations show that it can successfully disable cell growth upon burden-reducing mutations of different synthetic circuits’ genes, hindering the takeover of engineered cell population by mutants (Sections 2.1, 3.2 and 4). Importantly, the same DNA implementation of the Punisher can be reused across various applications simply by adjusting the culture medium’s chemical inducer content (Section 2.3), which avoids costly DNA re-design and synthesis steps characteristic of extant mutation spread mitigation strategies [6, 8]. Moreover, the Punisher directly senses resource competition through which synthetic gene expression impairs the growth of engineered cells. Consequently, it is only triggered when synthetic gene expression loss accelerates the mutant cell’s growth, whereas other approaches may exhibit unintended reactions to mutations that do not provide a growth advantage, inadvertently making engineered cell populations less genetically stable (Section 3.2).

Our design was studied using a coarse-grained resource-aware cell model [5], enabling a holistic view of cell growth’s burden-dependence that underlies mutation spread in cell populations, as well as the contribution of native and synthetic genes to the resource competition sensed by the Punisher, and the effect of antibiotics which (when resistance is disabled by our circuit) are used to penalise mutations. Possessing more predictive power than the basic gene expression models that ingore the cellular context, our modelling framework is also less complex than finer-grained cell models. This enables mathematical derivations that elucidate the Punisher’s switching behaviour and allow to determine the threshold value of burden at which it becomes activated. Furthermore, the computational efficiency of cell model simulations has enabled us to define the rates of cells switching between different states in a population based on stochastic single-cell trajectories rather than arbitrary parameterisation [7]. Given that synthetic genes in our model are described using physiologically relevant gene expression parameters like promoter strength and ribosome affinity, our approach establishes a novel rigorous method of directly linking a circuit’s (burden-dependent) population dynamics to its design parameters.

Whilst this study includes extensive simulations and analysis of the Punisher’s performance, important avenues of research remain to be explored. Our model currently assumes that the cell hosts just one copy of every burdensome synthetic gene whose mutations we aim to penalise, and that each gene is either fully functional or completely disabled by mutation, whereas in reality mutations may only partially reduce synthetic gene expression. Such partially-disabling mutations or mutations of just one among many synthetic gene copies may still be penalised by the Punisher if its switching threshold is carefully positioned between the burden experienced by the original engineered cells and the gene expression burden in such ‘incompletely’ mutated cells. However, simulating the engineered cell population’s behaviour in this case would require increasing the number of cell states considered, potentially making efficient implementation of such a model challenging. Another future direction of research into population modelling is the simulation of clonal interference dynamics for native gene mutations as speculated in Section 4.2. Finally, the Punisher circuit remains to be implemented and tested *in vivo*.

In conclusion, we have proposed a promising versatile biomolecular controller design, the same genetic implementation of which can be used to improve the genetic stability of engineered cell populations in a wide range of applications. More generally, the present study represents a showcase and a blueprint for how insights from cell modelling both on a single-cell and population level can be leveraged in several different aspects of resource-aware biomolecular controller development.

## Data and code accessibility

All data and code used in this publication are available at github.com/KSechkar/punisher.

## Author contributions

K.S.: Conceptualisation; Formal analysis; Investigation; Methodology; Software; Validation; Visualization; Writing – original draft; Writing – review & editing. H.S.: Conceptualisation; Funding acquisition; Project administration; Resources; Supervision; Writing – review & editing.

