## Supplementary Notes, Figures and Tables for "Model-guided gene circuit design for engineering genetically stable cell populations in diverse applications"

**S1 Cell and circuit modelling** **3**

### S1 Cell and circuit modelling

The main text describes the 'Punisher' circuit, which aims to detect mutation-induced alleviations of burden experienced by the cell hosting it and slow down the mutant host cell's growth to increase the genetic stability of engineered cell populations. In order to simulate and analyse our design's performance, we leverage a coarse-grained resource-aware cell model published in [1]. This framework consists of six ordinary differential equations (ODEs) that describe the expression of the native genes of an *Escherichia coli* cell. The expression of the synthetic genes hosted by this bacterium is then captured by extending the 'native' ODEs with additional equations in a standard form. Hence, specifying a modelled scenario of interest in this framework requires explicitly defining the added ODEs for synthetic genes and the parameters appearing in them. Moreover, the particularities of the Punisher's design means that several minor modifications must be made to the original cell model. These synthetic circuit ODEs and changes to the model, as well as the derivations associated with them, are provided in this supplementary note.

#### S1.1 Cell model ODEs

The coarse-grained cell model aims to capture the allocation of translational resources (i.e. the cell's ribosomes) between the synthesis of native and synthetic genes, as well as how this allocation influences cell growth [1]. To this end, the cell's genome is split into three classes by function and regulatory behaviour – ribosomal ($r$), metabolic ($a$) and housekeeping ($q$). The expression of housekeeping proteins, which always occupy a constant fraction $\overline{\phi}_q$ of the cell's overall protein mass $M$, is not modelled explicitly. Meanwhile, each of the former two classes are treated as a single lumped gene. The dynamics of these lumped genes' corresponding concentrations of mRNA ($m_r$ and $m_a$) and protein ($R$ and $p_a$, respectively) are modelled with ODEs. Similar equations are used to capture the dynamics of the mRNA and protein levels $m_{x_l}$ and $p_{x_l}$ for each gene $x_l$ from the set $X$ of synthetic genes hosted by the cell.

In order to maximise its growth rate in a given environment, the *E. coli* dynamically reallocates its resources by regulating the expression of its ribosomal genes. This is enabled by the ppGpp signalling molecule, whose concentration is proportional to the ratio of the charged (aminoacylated) and uncharged tRNA molecule abundances in the cell $t^c$ and $t^u$, hence the need to define ODEs for these variables as well [1]. Another factor determining translational resource availability is the potential presence of the ribosome-inhibiting antibiotic chloramphenicol, present in the cell at an intracellular concentration $h$. These considerations give rise to Equations (S1)–(S10), whose parameter and terms are specified and explained in Supplementary Table S1 and Supplementary Table S2, respectively.

$$\dot{m}_a = c_a\alpha_a\lambda(\epsilon, B) - (\beta_a + \lambda(\epsilon, B))m_a \tag{S1}$$

$$\dot{m}_r = F_r(t^u, t^c) \cdot c_r\alpha_r\lambda(\epsilon, B) - (\beta_r + \lambda(\epsilon, B))m_r \tag{S2}$$

$$\dot{p}_a = \frac{\epsilon(t^c)}{n_a} \cdot \frac{m_a/k_a}{D}R - \lambda(\epsilon, B) \cdot p_a \tag{S3}$$

$$\dot{R} = \frac{\epsilon(t^c)}{n_r} \cdot \frac{m_r/k_r}{D}R - \lambda(\epsilon, B) \cdot R \tag{S4}$$

$$\dot{t^c} = \nu(t^u, \sigma) \cdot p_a - \epsilon(t^c) \cdot B - \lambda(\epsilon, B) \cdot t^c \tag{S5}$$

$$\dot{t^u} = \psi(t^u, t^c) \cdot \lambda(\epsilon, B) - \nu(t^u, \sigma) \cdot p_a + \epsilon(t^c) \cdot B - \lambda(\epsilon, B) \cdot t^u \tag{S6}$$

$$\dot{h} = \kappa(h_{ext} - h) - \frac{hp_{cat}}{K_C} - \lambda(\epsilon, B) \cdot h \tag{S7}$$

$$\dot{m}_{x_l} = F_{x_l}(\cdot)c_{x_l}\alpha_{x_l}\lambda(\epsilon, B) - (\beta_{x_l} + \lambda(\epsilon, B))m_{x_l} \qquad \text{for all } x_l \in X \tag{S8}$$

$$\dot{p}_{x_l} = \frac{\epsilon(t^c)}{n_{x_l}} \cdot \frac{m_{x_l}/k_{x_l}}{D}R - (\delta_{x_l}p_{prot} + \lambda(\epsilon, B))p_{x_l} \qquad \text{for all } x_l \in X \tag{S9}$$

$$\text{where } D = \frac{K_D + h}{K_D} \cdot \left(1 + \frac{\sum_{j \in \{a,r\} \cup X} m_j/k_j - \frac{K_D+h}{K_D} \cdot \frac{\overline{\phi}_q \Delta}{\epsilon R}}{1 - \overline{\phi}_q \left(1 - \frac{K_D+h}{K_D} \cdot \frac{\Delta}{\epsilon R}\right)}\right) \tag{S10}$$

Due to the specifics of the Punisher circuit – namely, the fact that it includes the chloramphenicol acetyltransferase (CAT) protein, which degrades chloramphenicol in the cell, and a synthetic protease – we introduce two differences between the cell model above and the original in [1]. First, the possibility of chloramphenicol degradation by CAT means that we need to consider the dynamics of its intracellular concentration using Equation (S7), based on [2] and [3]. Moreover, since the original cell model operated only with extracellular chloramphenicol level, we now have to use a slightly altered definition of the 'resource competition denominator' $D$ in Equation (S10), which we derive in Supplementary Note S1.3. The dynamics of $p_{cat}$ and $p_{prot}$ appearing in this ODE are described by Equation (S18).

Second, synthetic protein degradation was formerly simplistically assumed to happen at a fixed rate with the help of the cell's own degradation machinery (for native protein levels, the effect of active degradation is negligible and does not need to be considered [1, 4]). Conversely, here synthetic proteins are broken down by the Punisher circuit's synthetic protease, whose availability it may be useful to adjust (see Supplementary Note S3.6). Hence, the degradation rate of synthetic proteins depends on its concentration $p_{prot}$ (see Equation (S17) for its dynamics).

Supplementary Table S1: **Parameters appearing in the cell model ODEs and Supplementary Table S2.**

| Parameter | Description | Value$^{\S}$ | Units$^{*}$ |
|---|---|---|---|
| $M$ | Total cell mass (in amino acids) | $1.19 \cdot 10^9$ | $aa^{\#}$ |
| $\sigma$ | Extracellular nutrient quality | $0.5^{\aleph}$ | None |
| $\overline{\phi}_q$ | Housekeeping prot. mass fraction$^{\S}$ | 0.59 | None |
| | **Reaction rates** | | |
| $\epsilon_{max}$ | Max. translation elongation rate | 72,000 | $aa^{\#}/h$ |
| $\nu_{max}$ | Max. tRNA charging rate | $4,046.9$ | $h^{-1}$ |
| $\psi_{max}$ | Max. tRNA synthesis per unit growth rate | $4.32 \cdot 10^5$ | $nM$ |
| $K_\epsilon$ | Michaelis constant for translation elongation | $1,239.7$ | $nM$ |
| $K_\nu$ | Michaelis constant for tRNA charging | $1,239.7$ | $nM$ |
| $\tau$ | Michaelis constant for ppGpp signalling | 1 | None |
| | **Gene expression** | | |
| $c_j$ | Concentration of gene $i$ DNA$^{\ddagger}$ | 1 | $nM$ |
| $\alpha_a$ | Promoter strength for gene $a$ | $3.945 \cdot 10^5$ | None |
| $\alpha_r$ | Promoter strength for gene $r$ | $4.070 \cdot 10^5$ | None |
| $\beta_i$ | mRNA degradation rate$^{\ddagger}$ | 6 | $h^{-1}$ |
| $k_j^+$ | mRNA-ribosome binding rate$^{\ddagger}$ | 60 | $\frac{1}{nM \cdot h}$ |
| $k_j^-$ | mRNA-ribosome dissociation rate$^{\ddagger}$ | 60 | $h^{-1}$ |
| $n_a$ | Number of amino acids in protein $p_a$ | 300 | $aa^{\#}/nM$ |
| $n_r$ | Number of amino acids in rib. protein | 7,459 | $aa^{\#}/nM$ |
| | **Chloramphenicol dynamics** | | |
| $h_{ext}$ | Chloramphenicol concentration in the culture medium | $10,500^{\aleph}$ | $nM$ |
| $\kappa$ | Chloramphenicol diffusion rate through the membrane | 5,400 [3] | $h^{-1}$ |
| $K_C$ | CAT-chloramphenicol affinity constant | 0.00556 [3] | $nM \cdot h$ |
| $K_D$ | Chloramphenicol-ribosome dissociation constant | 1,300 [3] | $nM$ |

$^{\S}$Same as for the original cell model in [1] unless stated otherwise.
$^{*}$*E. coli* volume is $\approx 10^{-18}$ m$^3$, so 1 nM is roughly equivalent to 1 molecule/cell [5].
$^{\aleph}$Culture conditions with high nutrient quality and strong ribosome inactivation by antibiotics [6].
$^{\#}$Amino acid residues. $^{\ddagger}$Same for any native gene $j \in \{a, r\}$.

Supplementary Table S2: **Functions and notations appearing in the host cell model ODEs.**

| Func./Not. | Description | Formula | Units* |
|---|---|---|---|
| $B$ | Number of translating ribosomes | $(\frac{K_D+h}{K_D} - \frac{1}{D}) \cdot R$ | $nM$ |
| $F_r$ | Transcription regulation for ribosomal genes (via ppGpp) | $\frac{t^c/t^u}{t^c/t^u+\tau}$ | None |
| $F_{x_l}(\cdot)$ | Transcription regulation for synthetic genes | 1 if $x_l$ constitutive, otherwise gene-specific | None |
| $\Delta$ | Total flux for protein degradation by the protease | $\sum_{x_l \in X} \delta_{x_l} p_{x_l} p_{prot}$ | $aa^{\#}/h$ |
| **Ribosome affinities** | | | |
| $k_i$ | mRNA-ribosome dissociation constant for gene $i^{\ddagger}$ | $\frac{k_i^- + \epsilon/n_i}{k_i^+}$ | $nM$ |
| **Reaction rates** | | | |
| $\epsilon$ | Translation elongation rate | $\epsilon_{max} \cdot \frac{t^c}{t^c+K_\epsilon}$ | $aa^{\#}/h$ |
| $\lambda$ | Growth/dilution rate | $\frac{\epsilon B}{M} - \frac{\Delta}{M}$ | $h^{-1}$ |
| $\psi$ | tRNA synthesis rate | $\psi_{max} \cdot \frac{t^c/t^u}{t^c/t^u+\tau}$ | $nM/h$ |
| $\nu$ | tRNA charging rate | $\nu_{max} \cdot \sigma \cdot \frac{t^u}{t^u+K_\nu}$ | $h^{-1}$ |

*$E. coli$ volume is $\approx 10^{-18}$ m$^3$, so 1 nM is roughly equivalent to 1 molecule/cell [5].
‡ Formula identical across all native **and** synthetic genes $i \in \{a, r\} \cup X$.
#Amino acid residues.

#### S1.2 Punisher circuit ODEs

In most cases considered in this study, the bacterial cell hosts the Punisher circuit, which includes the switch gene $s$, which activates its own expression when bound by a chemical inducer molecule, the synthetic protease *prot* degrading it, the chloramphenicol acetyltransferase (CAT) gene *cat* which confers antibiotic resistance by degrading chloramphenicol in the cell, and the integrase $i$ that can excise the CAT gene from the plasmid hosting it. Since the integrase is expressed from the same operon as the switch gene, their corresponding mRNA concnetrations are identical up to a scaling factor $^{n_i}/_{n_s}$, which arises because our model scales the mRNA concentration of a given gene $j$ by $^{n_j}/_{25}$ to account for the possibility of the same protein coding sequence being translated by mutliple ribosomes at a time [1]. The ODEs for the expression of these four genes $\{s,\ i,\ prot,\ cat\} \subseteq X$ are given by Equations (S11)–(S18). All synthetic gene parameters for this circuit are provided in Supplementary Table S3.

$$\dot{m}_s = F_s(p_s, I)c_s\alpha_s\lambda(\epsilon, B) - (\beta_s + \lambda(\epsilon, B))m_s \tag{S11}$$

$$m_i \equiv m_s \cdot \frac{n_i}{n_s} \text{ as } s \text{ and } i \text{ genes are co-expressed from the same operon} \tag{S12}$$

$$\dot{m}_{prot} = c_{prot}\alpha_{prot}\lambda(\epsilon, B) - (\beta_{prot} + \lambda(\epsilon, B))m_{prot} \tag{S13}$$

$$\dot{m}_{cat} = c_{cat}\alpha_{cat}\lambda(\epsilon, B) - (\beta_{cat} + \lambda(\epsilon, B))m_{cat} \tag{S14}$$

$$\dot{p}_s = \frac{\epsilon(t^c)}{n_s} \cdot \frac{m_s/k_s}{D}R - (\delta_s p_{prot} + \lambda(\epsilon, B))p_s \tag{S15}$$

$$\dot{p}_i = \frac{\epsilon(t^c)}{n_i} \cdot \frac{m_i/k_i}{D}R - \lambda(\epsilon, B) \cdot p_i - \delta_i p_i \tag{S16}$$

$$\dot{p}_{prot} = \frac{\epsilon(t^c)}{n_{prot}} \cdot \frac{m_{prot}/k_{prot}}{D}R - \lambda(\epsilon, B) \cdot p_{prot} \tag{S17}$$

$$\dot{p}_{cat} = \frac{\epsilon(t^c)}{n_{cat}} \cdot \frac{m_{cat}/k_{cat}}{D}R - \lambda(\epsilon, B) \cdot p_{cat} \tag{S18}$$

$$\text{where } F_s(p_s, I) = F_{s_b} + (1 - F_{s_b}) \cdot \frac{(Ip_s)^{\eta_s}}{(Ip_s)^{\eta_s} + K_s^{\eta_s}} \tag{S19}$$

While the DNA concentrations of most genes are constant parameters, the CAT gene can cut out of the plasmid and become non-functional due to integrase action. We therefore use Equations (S20) and (S21) to respectively model the dynamics of the concentration of functional CAT gene DNA $c_{cat}$ and the abundance $c_{LRi}$ of strand exchange products formed in the first step of gene excision by the integrase. These ODEs are parameterised in Supplementary Table S3 and explained in Suppplementary Note S1.4. Assuming that the CAT gene is situated on the same synthetic plasmid as the rest of the Punisher circuit, we use the initial condition $c_{LRi}(t = 0\ h) = 0\ nM$ and $c_{cat}(t = 0\ h) = c_s = c_i = c_{prot} = 10\ nM$.

$$\dot{c}_{cat} = -k_{sx}^{+}\frac{p_i^4}{K_{bI}^4 + p_i^4}c_{cat} + k_{sx}^{-}c_{LRi} \tag{S20}$$

$$\dot{c}_{LRi} = k_{sx}^{+}\frac{p_i^4}{K_{bI}^4 + p_i^4}c_{cat} - k_{sx}^{-}c_{LRi} - (k^{conf} + \lambda)c_{LRi} \tag{S21}$$

Supplementary Table S3: Parameter values for simulating the Punisher circuit.

| Parameter | Description | Value[§] | Units[*] |
|---|---|---|---|
| $c_s = c_{prot}$ | Switch/integrase and protease gene DNA concentration | 10 | $nM$ |
| $\alpha_s$ | Switch gene promoter strength | 400 | None |
| $\alpha_{prot}$ | Protease gene promoter strength | 25 | None |
| $\alpha_{cat}$ | *cat* gene promoter strength | 500 | None |
| $\beta_j$ | mRNA degradation rates | 6 | $h^{-1}$ |
| $k_s^+ = k_{prot}^+ = k_{cat}^+$ | mRNA-ribosome binding rate for $s$, $prot$ and $cat$ genes | 60[ℵ] | $\frac{1}{nM \cdot h}$ |
| $k_i^+$ | mRNA-ribosome binding rate for the integrase gene | 0.75[ℵ] | $\frac{1}{nM \cdot h}$ |
| $k_s^- = k_i^- = k_{prot}^- = k_{cat}^-$ | mRNA-ribosome dissociation rate | 60 | $h^{-1}$ |
| $n_s = n_i = n_{prot} = n_{cat}$ | Number of amino acids in proteins | 300 | $aa/nM$ |
| $\delta_s$ | Rate of switch protein degradation by the protease | 0.01836 [7] | $\frac{1}{nM \cdot h}$ |
| $\delta_i = \delta_{prot} = \delta_{cat}$ | Rate of non-switch protein degradation by the protease (zero as no degradation tag) | 0 | $\frac{1}{nM \cdot h}$ |
| | **Gene transcription regulation functions** | | |
| $F_{s_b}$ | Baseline function value in absence of the inducer | 0.025 | None |
| $K_s$ | Half-saturation constant for for the binding between the switch gene promoter DNA and switch protein-inducer complexes | 300 | $nM$ |
| $\eta_s$ | Cooperativity coefficient for the binding between the switch gene promoter DNA and switch protein-inducer complexes | 2 | $nM$ |
| | **Integrase action** | | |
| $k_{sx}^+$ | Forward DNA strand exchange rate | 6 [8] | $h^{-1}$ |
| $k_{sx}^-$ | Reverse DNA strand exchange rate | 2.14 [8] | $h^{-1}$ |
| $k^{conf}$ | Rate of strand exchange product-integrase complex conformation change | 0.006 [8] | $h^{-1}$ |
| $K_{bI}$ | Dissociation constant for CAT gene DNA-integrase complex formation | 100 [8] | $nM$ |
| $I$ | Fraction of switch proteins bound by a chemical inducer molecule | Varies | None |

§Unless stated otherwise, picked from the biologically realistic synthetic gene parameter ranges estimated for the cell model in [1].

**E. coli* volume is $\approx 10^{-18}$ m$^3$, so 1 nM is roughly equivalent to 1 molecule/cell [5].

ℵFor genes other than the integrase, near the diffusion limit (strong RBS) [6]. For the integrase, 80 times weaker. Experimentally, an up to $\approx$ 250-fold difference between the strongest and the weakest RBS activity in a synthetic lirary has been observed [9, 10].

‡Same for any Punisher gene $j \in \{s,\ i,\ prot,\ cat\}$.

#### S1.3 Modelling chloramphenicol action

In this supplementary note, we derive the relations allowing our cell model to capture how the bacterial cell is affected by the intracellular concentration of the chloramphenicol antibiotic, which reversibly binds the ribosome's 50S subunit, which makes translating ribosomes stall on their transcripts [2, 3].

To this end, we start with a cell model that explicitly considers binding between the ribosomes and the mRNA and between the ribosomes and chloramphenicol, as well as the expression of the cell's housekeeping genes (unlike our main cell model in Equations (S1)–(S10), where housekeeping gene expression is considered implicitly). Analogically to the derivations in [1], we then reduce this extended cell model to the simplified version in Equations (S1)–(S10).

The newly added explicit considerations mean that besides those discussed in Supplementary Note S1.1, there are now more variables to be modelled, such as housekeeping mRNA and protein concentrations $m_q$ and $p_q$. Moreover, instead of modelling the dynamics of the ribosomes' overall abundance, we need to consider the concentration of free (i.e. not bound by any transcript) ribosomes $r$, chloramphenicol-bound free ribosomes $r^{cm}$, mRNA-ribosome complex concentrations $\{b_j\}$ for each gene $j \in \{q, a, r\} \cup X$, and chloramphenicol-bound (inactive) mRNA-ribosome complex abundances $\{b_j^{cm}\}$. This gives rise to the Equations (S22)–(S31). The transcription of housekeeping genes here obeys the transcription regulation function $F_q$ (which we do not specify here), while all parameters of housekeeping genes are assumed to be the same as for the metabolic gene class $a$ [1]. Also note that $k_D^+$ and $k_D^-$ are the forward and reverse binding rates between the ribosome and a chloramphenicol molecule. All other parameters and notations can be found in Supplementary Tables S1–S2.

$$\dot{m_j} = F_j c_j \alpha_j \lambda(\epsilon, B) - (\beta_j + \lambda(\epsilon, B))m_j - k_j^+ m_j r + k_j^- b_j + \frac{\epsilon(t^c)}{n_j} b_j \quad \text{for } j \in \{q, a, r\} \cup X \quad \text{(S22)}$$

$$\dot{b_j} = k_j^+ m_j r - k_j^- b_j - \frac{\epsilon(t^c)}{n_j} b_j - \lambda(\epsilon, B) \cdot b_j - k_D^+ h b_j + k_D^- b_j^{cm} \quad \text{for } j \in \{q, a, r\} \cup X \quad \text{(S23)}$$

$$\dot{b_j^{cm}} = k_D^+ h b_i - k_D^- b_j^{cm} - \lambda(\epsilon, B) \cdot b_j^{cm} \quad \text{for } j \in \{q, a, r\} \cup X \quad \text{(S24)}$$

$$\dot{p_j} = \frac{\epsilon(t^c)}{n_j} b_j - \lambda(\epsilon, B) \cdot p_j \quad \text{for } j \in \{q, a\} \quad \text{(S25)}$$

$$\dot{p_{x_l}} = \frac{\epsilon(t^c)}{n_{x_l}} b_{x_l} - (\lambda(\epsilon, B) + \delta_{x_l} p_{prot}) p_{x_l} \quad \text{for } x_l \in X \quad \text{(S26)}$$

$$\dot{r} = \frac{\epsilon(t^c)}{n_r} b_r + \sum_{j \in \{q,a,r\} \cup X} \left( (\frac{\epsilon(t^c)}{n_j} + k_j^-) b_j - k_j^+ m_j r \right) - k_D^+ h r + k_D^- r^{cm} - \lambda(\epsilon, B) \cdot r \tag{S27}$$

$$\dot{r}^{cm} = -k_D^+ h r + k_D^- r^{cm} - \lambda(\epsilon, B) \cdot r^{cm} \tag{S28}$$

$$\dot{t}^c = \nu(t^u, \sigma) \cdot p_a - \epsilon(t^c) \cdot B - \lambda(\epsilon, B) \cdot t^c \tag{S29}$$

$$\dot{t}^u = \psi(T) \cdot \lambda(\epsilon, B) - \nu(t^u, \sigma) \cdot p_a + \epsilon(t^c) \cdot B - \lambda(\epsilon, B) \cdot t^u \tag{S30}$$

$$\dot{h} = \kappa(h_{ext} - h) - \frac{h p_{cat}}{K_C} - \lambda(\epsilon, B) \cdot h \tag{S31}$$

Using the fact that binding and unbinding between mRNAs and ribosomes, as well as between ribosomes and chloramphenicol, happen at a much faster timescale than all other cellular processes in our model [1, 2], we assume that the concentrations of mRNA-ribosome and ribosome-chloramphenicol complexes are in quasi-steady state. Hence,

$$b_j = \frac{m_j}{k_j} r \quad \text{for} \quad j \in \{q, a, r\} \cup X \tag{S32}$$

$$\text{and} \quad b_j^{cm} = \frac{h}{K_D} b_j, \quad r^{cm} = \frac{h}{K_D} r \tag{S33}$$

where

$$k_j = \frac{k_j^- + \epsilon(t^c)/n_j + \lambda}{k_j^+} \approx \frac{k_j^- + \epsilon/n_i}{k_j^+} \tag{S34}$$

$$\text{and} \quad K_D = \frac{k_D^- + \lambda}{k_D^+} \approx \frac{k_D^-}{k_D^+} \tag{S35}$$

are the dissociation constants for ribosome binding to mRNAs and chloramphenicol, respectively. The approximations allowing to neglect the contribution of the cell growth rate are based on the fact that $(k_j^- + \epsilon/n_j) \gg \lambda$ [1] and $k_D^- \gg \lambda$ [2]. The total concentration of ribosomes is therefore given by Equation (S36), and the synthesis rate for $r$ and $p_j$ $\forall j \in \{q, a\} \cup X$ is yielded by Equation (S37).

$$R = r + r^{cm} + \sum_{j \in \{q,a,r\} \cup X} (b_j + b_j^{cm}) = \frac{K_D + h}{K_D} \cdot \left( 1 + \sum_{j \in \{q,a,r\} \cup X} \frac{m_j}{k_j} \right) \cdot r \tag{S36}$$

$$\frac{\epsilon}{n_j} \cdot b_j = \frac{\epsilon}{n_j} \cdot \frac{m_j/k_j}{\frac{K_D+h}{K_D} \cdot \left( 1 + \sum_{l \in \{q,a,r\} \cup X} \frac{m_l}{k_l} \right)} \cdot R \tag{S37}$$

Equation (S37) allows to avoid explicitly considering the concentrations of free ribosomes and mRNA-ribosome complexes, instead expressing the rates of protein synthesis in terms of the total ribosome abundance $R$ and mRNA concentrations $\{m_j\}$. However, it still includes the housekeeping mRNA concentration $m_q$. To get rid of it, we recall that the regulation of housekeeping protein expression $F_q$ is such that the mass fraction of housekeeping proteins in the cell's overall proteome is always $\overline{\phi}_q = 0.59$ [1]. This means that the housekeeping protein concentration is in a steady state. Solving $\dot{p}_q = 0$, we get:

$$\frac{m_q}{k_q} = \frac{\overline{\phi}_q \sum_{j \in \{a,r\} \cup X} m_j/k_j - \overline{\phi}_q \cdot \frac{K_D + h}{K_D} \cdot (1 + \sum_{j \in \{a,r\} \cup X} m_j/k_j) \cdot \frac{\Delta}{\epsilon R}}{1 - \overline{\phi}_q \left(1 - \frac{K_D + h}{K_D} \cdot \frac{\Delta}{\epsilon R}\right)} \tag{S38}$$

Substituting Equation (S38) into Equation (S37) yields Equation (S39), where the formula for $D$ matches Equation (S10) from our main cell model described in Supplementary Note S1.1.

$$\frac{\epsilon}{n_j} \cdot b_j = \frac{\epsilon}{n_j} \cdot \frac{m_j/k_j}{\frac{K_D + h}{K_D} \cdot \left(1 + \frac{\sum_{j \in \{a,r\} \cup X} m_j/k_j - \frac{K_D+h}{K_D} \cdot \frac{\overline{\phi}_q \Delta}{\epsilon R}}{1 - \overline{\phi}_q \left(1 - \frac{K_D+h}{K_D} \cdot \frac{\Delta}{\epsilon R}\right)}\right)} \cdot R = \frac{\epsilon}{n_j} \cdot \frac{m_j/k_j}{D} \cdot R \tag{S39}$$

Let us now consider once again the extended cell model in (S22)–(S31). Equations (S32)–(S37) mean that we no longer need to individually consider the concentrations of ribosomes that are bound by mRNAs, chloramphenicol, both, or neither, since protein synthesis rates can now be defined using the total ribosome abundance $R$. The ODE for $R = r + r^{cm} + \sum_{j \in \{q,a,r\} \cup X} (b_j + b_j^{cm})$ can be obtained by adding together the ODEs for the abundances of ribosomes in all the states that we used to treat individually. Moreover, due to our quasi-steady-state assumption, the rates of ribosome-mRNA binding and unbinding are equal, so in the mRNA concentration ODE (S22) they cancel each other out. Finally, Equation (S39) allows to get rid of the ODEs that describe the dynamics of housekeeping protein and mRNA concentrations. These simplifications transform our extended cell model into our main cell model in Supplementary Note S1.1's Equations (S1)–(S10).

In summary, we have derived our main cell model in Equations (S1)–(S10) starting from an extended mechanistic model in (S22)–(S31), which explicitly incorporated the binding and unbinding between the cell's ribosomes and chloramphenicol molecules present in the cell. This demonstrates that our main cell model indeed captures the effects of chloramphenicol's intracellular concentration on the host cell's functioning.

#### S1.4 Modelling integrase action

The Punisher circuit described in this study involves a serine integrase, which is capable of excising a gene essential for cell growth (e.g. the antibiotic resistance gene CAT) from its place in a synthetic plasmid or the host cell's genome. When synthetic gene mutation reduces the competition for ribosomes in the cell, integrase expression is upregulated and the essential gene is excised, becoming unable to be expressed. This loss of essential gene expression underlies the penalisation of mutations enacted by the Punisher and thus is a vital part of our design's functioning that must be modelled appropriately, which we do using Equations (S20)–(S21). The present supplementary note outlines the considerations behind these ODEs, which are primarily based on the serine integrase-mediated DNA recombination model in [11] and [8].

Originally, the gene of interest on the plasmid is flanked by the integrase's cognate sites attP and attB. Its excision can then be said to occur in three steps [11]. First, the attP and attB sites on the plasmid's DNA are bound by four integrase molecules (two per site). Second, the integrase tetramer exchanges the DNA strands to produce attL and attR sites – one remaining in the plasmid, another in the newly formed small circular DNA fragment containing the CAT gene – which are still held together by the tetramer. Finally, the complex changes its conformation or is broken apart by the protein machinery that replicates the plasmid to keeps its copy number constant despite dilution due to cell growth. The first two steps are reversible but the last one is not, since the excised gene now becomes a circular DNA fragment which diffuses away from its original location on the plasmid. The whole process can thus be represented as:

$$attPattB + 4p_i \underset{k^+_{bind}}{\overset{k^-_{bind}}{\rightleftarrows}} attPattB:(p_i)_4 \underset{k^+_{sx}}{\overset{k^-_{sx}}{\rightleftarrows}} attLattR:(p_i)_4 \xrightarrow{k^{conf}+\lambda} attL + attR + 4p_i$$

Since the binding and unbinding between the attPattB sites and the integrase are much faster than other modelled processes, we assume that the concentration of the $attPattB:(p_i)_4$ complex is in a quasi-steady state. Assuming that the fraction of the DNA-bound integrase is negligible compared to that of the free protein [11], the concentration of $attPattB:(p_i)_4$ can be found from the total concetration of the CAT gene's non-excised DNA $c_{cat}$ as:

$$\frac{p_i^4}{K_{bI}^4 + p_i^4} c_{cat} \tag{S40}$$

This allows us to only model the total functional CAT gene concentration and the concentration $c_{LRi}$ of the integrase-attLattR site complexes that form as a result of reversible DNA strand exchange. Their dynamics are thus described by Equations (S20)–(S21), that is,

$$\dot{c}_{cat} = -k_{sx}^+ \frac{p_i^4}{K_{bI}^4 + p_i^4} c_{cat} + k_{sx}^- c_{LRi}$$
$$\dot{c}_{LRi} = k_{sx}^+ \frac{p_i^4}{K_{bI}^4 + p_i^4} c_{cat} - k_{sx}^- c_{LRi} - (k^{conf} + \lambda) c_{LRi}$$

While the derived ODEs can be used for deterministic simulations, we also perform stochastic simulations of the Punisher's behaviour, performed according to the hybrid tau-leaping algorithm described in [1]. In this case, $c_{cat}$ and $c_{LRi}$ are discrete random non-negative integer variables representing the numbers of gene copies in the cell (since $1\ nM \approx 1\ molecule/cell$ [5], we use the initial condition $c_{LRi} = 0$ and $c_{cat} = 10$). In order to simulate their evolution over time, the following reactions are added to the standard stochastic gene expression processes:

- **Forward DNA strand exchange** at rate $k_{sx}^+ \frac{p_i^4}{K_{bI}^4 + p_i^4} c_{cat}$. Decreases $c_{cat}$ by 1, increases $c_{LRi}$ by 1

- **Reverse DNA strand exchange** at rate $k_{sx}^- c_{LRi}$. Increases $c_{cat}$ by 1, decreases $c_{LRi}$ by 1

- **Synaptic conformations change of the integrase-attLattR site complex** at rate $k^{conf} c_{LRi}$. Decreases $c_{LRi}$ by 1

- **Breakdown of the integrase-attLattR site complex by plasmid replication machinery** at rate $\lambda c_{LRi}$. Decreases $c_{LRi}$ by 1

#### S1.5 Synthetic gene circuit ODEs

The Punisher circuit described in our study aims to detect and penalise changes in gene expression burden that arise due to mutations in other synthetic gene circuitry also present in the cell. This supplementary note provides the explicit ODE definitions for these additional synthetic circuits, as well as parameterises them. Moreover, in order to model an alternative to our design and compare the Punisher to it, in Section S1.5.3 we provide the ODEs and parameters for a synthetic circuit whose mutations are penalised through the essential gene co-expression mechanism rather than our Punisher design.

##### S1.5.1 Single synthetic burdensome gene

In the simplest case, considered in the main text's Section 2.1 and Figure 1, a single constitutive burdensome synthetic gene $b$ is expressed in addition to the Punisher circuit. The ODEs for its mRNA and protein concentrations $m_b$ and $p_b$ are obtained straightforwardly from the generalised cell model Equations (S8)–(S9), which yields Equations (S41)–(S42). Due to the gene being constitutive, its transcription regulation function is always $F_b \equiv 1$. All synthetic gene expression parameters are provided in Supplementary Table S4. Whenever we simulate losing gene $b$ expression due to mutation, this is achieved by setting its concentration to $c_b = 0\ nM$.

$$\dot{m}_b = F_b \cdot c_b \alpha_b \lambda(\epsilon, B) - (\beta_b + \lambda(\epsilon, B)) m_b \tag{S41}$$

$$\dot{p}_b = \frac{\epsilon(t^c)}{n_b} \cdot \frac{m_b/k_b}{D} R - (\delta_b p_{prot} + \lambda(\epsilon, B)) p_b \tag{S42}$$

Supplementary Table S4: Gene $b$ parameters.

| Parameter | Description | Value§ | Units* |
|---|---|---|---|
| $c_b$ | Gene copy number | 1 | $nM$ |
| $\alpha_b$ | Promoter strength | 100,000 [12] | None |
| $\beta_b$ | mRNA degradation rate | 6 | $h^{-1}$ |
| $k_b^+$ | mRNA-ribosome binding rate | 60 | $\frac{1}{nM \cdot h}$ |
| $k_b^-$ | mRNA-ribosome dissociation rates | 60 | $h^{-1}$ |
| $n_b$ | Number of amino acids in protein | 300 | $aa/nM$ |
| $\delta_b$ | Rate of protein degradation by the Punisher's protease (zero as no degradation tag) | 0 | $\frac{1}{nM \cdot h}$ |

§Unless stated otherwise, picked from the biologically realistic synthetic gene parameter ranges estimated for the cell model in [1].
*E. coli volume is $\approx 10^{-18}$ m$^3$, so 1 nM is roughly equivalent to 1 molecule/cell [5].

##### S1.5.2 Two synthetic toggle switches

In the main text's Section 3.1 and Figure 3, we consider the case of a more complex synthetic gene circuit being present in the cell alongside the Punisher. Namely, the cell is conferred with two toggle switch circuits. A toggle switch consists of a pair of genes, each of which encodes a transcription factor that represses the other gene's expression. The strength of this repression can be modulated by the concentration of a chemical inducer in the culture medium, whose molecules may form complexes with the transcription factors and prevent them from binding the target promoter's DNA. Denoting the first toggle's genes as $tog_{11}$ and $tog_{12}$, and the second switch's genes as $tog_{21}$ and $tog_{22}$, we obtain the ODEs given by Equations (S43)–(S50).

$$\dot{m}_{tog_{11}} = F_{tog_{11}}(p_{tog_{12}}, I_{11}) \cdot c_{tog_{11}}\alpha_{tog_{11}}\lambda(\epsilon, B) - (\beta_{tog_{11}} + \lambda(\epsilon, B))m_{tog_{11}} \tag{S43}$$

$$\dot{m}_{tog_{12}} = F_{tog_{12}}(p_{tog_{11}}, I_{12}) \cdot c_{tog_{12}}\alpha_{tog_{12}}\lambda(\epsilon, B) - (\beta_{tog_{12}} + \lambda(\epsilon, B))m_{tog_{12}} \tag{S44}$$

$$\dot{m}_{tog_{21}} = F_{tog_{21}}(p_{tog_{22}}, I_{21}) \cdot c_{tog_{21}}\alpha_{tog_{21}}\lambda(\epsilon, B) - (\beta_{tog_{21}} + \lambda(\epsilon, B))m_{tog_{21}} \tag{S45}$$

$$\dot{m}_{tog_{22}} = F_{tog_{22}}(p_{tog_{21}}, I_{22}) \cdot c_{tog_{22}}\alpha_{tog_{22}}\lambda(\epsilon, B) - (\beta_{tog_{22}} + \lambda(\epsilon, B))m_{tog_{22}} \tag{S46}$$

$$\dot{p}_{tog_{11}} = \frac{\epsilon(t^c)}{n_{tog_{11}}} \cdot \frac{m_{tog_{11}}/k_{tog_{11}}}{D} R - (\delta_{tog_{11}} p_{prot} + \lambda(\epsilon, B))p_{tog_{11}} \tag{S47}$$

$$\dot{p}_{tog_{12}} = \frac{\epsilon(t^c)}{n_{tog_{12}}} \cdot \frac{m_{tog_{12}}/k_{tog_{12}}}{D} R - (\delta_{tog_{12}} p_{prot} + \lambda(\epsilon, B))p_{tog_{12}} \tag{S48}$$

$$\dot{p}_{tog_{21}} = \frac{\epsilon(t^c)}{n_{tog_{21}}} \cdot \frac{m_{tog_{21}}/k_{tog_{21}}}{D} R - (\delta_{tog_{21}} p_{prot} + \lambda(\epsilon, B))p_{tog_{21}} \tag{S49}$$

$$\dot{p}_{tog_{22}} = \frac{\epsilon(t^c)}{n_{tog_{22}}} \cdot \frac{m_{tog_{22}}/k_{tog_{22}}}{D} R - (\delta_{tog_{22}} p_{prot} + \lambda(\epsilon, B))p_{tog_{22}} \tag{S50}$$

The toggle genes' transcription regulation functions in the above ODEs are given by:

$$F_{tog_{11}}(p_{tog_{12}}, I_{11}) = F_{tog_{11},b} + (1 - F_{tog_{11},b}) \cdot \frac{K_{tog_{11}}^{\eta_{tog_{11}}}}{((1 - I_{11})p_{tog_{12}})^{\eta_{tog_{11}}} + K_{tog_{11}}^{\eta_{tog_{11}}}} \tag{S51}$$

$$F_{tog_{12}}(p_{tog_{11}}, I_{12}) = F_{tog_{12},b} + (1 - F_{tog_{12},b}) \cdot \frac{K_{tog_{12}}^{\eta_{tog_{12}}}}{(1 - I_{12})p_{tog_{11}})^{\eta_{tog_{12}}} + K_{tog_{12}}^{\eta_{tog_{12}}}} \tag{S52}$$

$$F_{tog_{21}}(p_{tog_{22}}, I_{21}) = F_{tog_{21},b} + (1 - F_{tog_{21},b}) \cdot \frac{K_{tog_{21}}^{\eta_{tog_{21}}}}{((1 - I_{21})p_{tog_{22}})^{\eta_{tog_{21}}} + K_{tog_{21}}^{\eta_{tog_{21}}}} \tag{S53}$$

$$F_{tog_{22}}(p_{tog_{21}}, I_{22}) = F_{tog_{22},b} + (1 - F_{tog_{22},b}) \cdot \frac{K_{tog_{22}}^{\eta_{tog_{22}}}}{(1 - I_{22})p_{tog_{21}})^{\eta_{tog_{22}}} + K_{tog_{22}}^{\eta_{tog_{22}}}} \tag{S54}$$

For simplicity, we assume that all toggle genes have identical expression dynamics (up to the indentity of the genes repressing their transcription as per Equations (S51)–(S54), so in Table S5 their values are given for a single gene $tog_{yz}$ with the implication that they are the same across $tog_{11}$, $tog_{12}$, $tog_{21}$ and $tog_{22}$. By default, no toggle gene inducers are assumed to be present in the culture medium, i.e.

$$I_{11} = I_{12} = I_{21} = I_{22} = 0 \tag{S55}$$

In order to break the symmetry and make sure that initially the two toggles have highly expressed $tog_{11}$ and $tog_{21}$ genes respectively, we set the initial condition:

$$m_{tog_{11}}(t = 0\ h) = m_{tog_{21}}(0\ h) = 4,000\ nM, \quad m_{tog_{12}}(t = 0\ h) = m_{tog_{22}}(0\ h) = 0\ nM \tag{S56}$$

$$p_{tog_{11}}(t = 0\ h) = p_{tog_{12}}(0\ h) = p_{tog_{21}}(t = 0\ h) = p_{tog_{22}}(0\ h) = 0\ nM \tag{S57}$$

The flipping of toggle 1 from high $tog_{11}$ expression to high $tog_{12}$ expression via an inducer concentration pulse was simulated by setting $I_{12} = 1$ for 45 minutes (we then reset this value to 0 as per Equation (S55)). The flipping of the second toggle to high $tog_{22}$ expression was simulated analogically. Likewise to the single constitutive gene in Supplementary Note S1.5.1, mutation of a given gene $tog_{yz}$ was simulated by setting $c_{tog_{yz}} = 0\ nM$.

Supplementary Table S5: Toggle switch gene parameters (identical for all four genes, hence the generic index $yz$).

| Parameter | Description | Value§ | Units* |
|---|---|---|---|
| $c_{tog_{yz}}$ | Gene copy number | 1 | $nM$ |
| $\alpha_{tog_{yz}}$ | Promoter strength | 50,000ℵ | None |
| $\beta_{tog_{yz}}$ | mRNA degradation rate | 6 | $h^{-1}$ |
| $k^{+}_{tog_{yz}}$ | mRNA-ribosome binding rate | 60 | $\frac{1}{nM \cdot h}$ |
| $k^{-}_{tog_{yz}}$ | mRNA-ribosome dissociation rates | 60 | $h^{-1}$ |
| $n_{tog_{yz}}$ | Number of amino acids in protein | 300 | $aa/nM$ |
| $\delta_{tog_{yz}}$ | Rate of protein degradation by the Punisher's protease (zero as no degradation tag) | 0 | $\frac{1}{nM \cdot h}$ |
| | **Transcription regulation function** | | |
| $F_{tog_{yz},b}$ | Baseline function value in absence of inducer | 0.025 | None |
| $K_{tog_{yz}}$ | Half-saturation constant for the repressor-promoter binding | 2,500 | $nM$ |
| $\eta_{tog_{yz}}$ | Cooperativity coefficient for the repressor-promoter binding | 2 | $nM$ |
| $I_{yz}$ | Fraction of proteins repressing $tog_{yz}$ gene transcription rendered unable to do so because of being bound by a chemical inducer molecule | Varied | None |

§Unless stated otherwise, picked from the biologically realistic synthetic gene parameter ranges estimated for the cell model in [1].
*E. coli volume is $\approx 10^{-18}$ m$^3$, so 1 nM is roughly equivalent to 1 molecule/cell [5].
ℵChosen to make the burden of fully expressing a single toggle's both genes equal to that imposed by a single burdensome gene in Supplementary Note S1.5.1.

##### S1.5.3 Two synthetic toggle switches with essential gene co-expression

In the main text's Section 3.4 and Figure 4, we compare the Punisher's performance with an extant essential gene co-expression strategy for countering mutation spread. Our Punisher design has a gene essential to cell growth that is expressed on its own but is susceptible to excision by the integrase if synthetic gene mutations are detected. Conversely, the co-expression approach involves coupling the expression of an essential gene with that of synthetic circuit genes – hence, mutations that disable synthetic gene expression also lower essential protein levels, which penalises the mutant cell by reducing its growth rate.

We apply the co-expression approach to the set of two synthetic toggle switches described in Supplementary Note S1.5.2. For simplicity, we assume that likewise to the Punisher, the essential gene used is the chloramphenicol acetyltransferase CAT, which confers antibiotic resistance to the cell. Hence, each of the toggle genes $\{tog_{11},\ tog_{12},\ tog_{21},\ tog_{22}\}$ is expressed in an operon with a CAT gene copy. The overall CAT protein synthesis rate is thus the sum of four translation rates as per Equations (S58)–(S59). Here, the index $cat_{yz}$ stands for the CAT gene copy co-expressed with the toggle gene $tog_{yz}$, $k_{cat_{yz}}$ is the mRNA-ribosome dissociation constant for the RBS of the corresponding CAT gene copy, calculated according to the formula in Supplementary Table S2 using the parameters from Supplementary Table S6. Likewise to Equation (S12) for the integrase's co-expression with the switch protein in the Punisher circuit, in Equation (S59) the toggle switch gene mRNA concentrations are scaled in order to be converted to the corresponding CAT mRNA concentrations, which is needed to account for translation of one transcript by several ribosomes at the same time.

$$\dot{p}_{cat} = \frac{\epsilon(t^c)}{n_{cat}} \cdot \frac{{m_{tog_{11}}}/{k_{cat_{11}}} + {m_{tog_{12}}}/{k_{cat_{12}}} + {m_{tog_{21}}}/{k_{cat_{21}}} + {m_{tog_{22}}}/{k_{cat_{22}}}}{D} R - \lambda(\epsilon, B) \cdot p_{cat} \tag{S58}$$

$$\text{where} \quad m_{cat_{yz}} \equiv m_{tog_{yz}} \cdot \frac{n_{cat}}{n_{tog_{yz}}} \quad \forall yz \in \{11,\ 12,\ 21,\ 22\} \tag{S59}$$

Note that the Punisher circuit is naturally absent from the cell if the co-expression method is used instead. Therefore, simulating two toggle switches with CAT co-expression involves concatenating the host cell model in Equations (S1)–(S7) **only** with Equations (S43)–(S54) for the toggle switches and Equation (S58) for the co-expressed CAT protein, but **not** the Punisher's ODEs. Since the Punisher's synthetic protease is also absent ($p_{prot} \equiv 0\ nM$), the degradation terms in all synthetic protein concentration ODEs are all zero, so the removal of proteins only happens via dilution due to cell division at the rate $\lambda$.

Besides comparing different mutation spread mitigation strategies, we benchmark them against the case of not using any strategies for hindering mutant cell growth. In this scenario, the two toggle switches are the only piece of synthetic circuitry present in the cell and there is no need for chloramphenicol to be present in the culture medium ($h_{ext} \equiv 0\ nM$, $p_{cat} \equiv 0\ nM$). Hence, the host cell model in Equations (S1)–(S7) is appended only with toggle switch ODEs (S43)–(S54).

Supplementary Table S6: CAT RBS parameters in the essential gene co-expression scenario (identical for all four CAT gene copies, hence the generic index $yz$).

| Func./Not. | Description | Value§ | Units* |
|---|---|---|---|
| $k^{+}_{cat_{yz}}$ | mRNA-ribosome binding rate | Varied | $\frac{1}{nM \cdot h}$ |
| $k^{-}_{cat_{yz}}$ | mRNA-ribosome dissociation rates | 60 | $h^{-1}$ |
| $n_{cat}$ | Number of amino acids in protein* | 300 | $aa/nM$ |

§Unless stated otherwise, picked from the biologically realistic synthetic gene parameter ranges estimated for the cell model in [1].

*E. coli* volume is $\approx 10^{-18}$ m$^3$, so 1 nM is roughly equivalent to 1 molecule/cell [5].

### S2 Additional circuit simulations

In this supplementary note, we display the outcomes of simulations that were mentioned in the main text but did not appear in the main text's figures for the sake of brevity and clarity.

#### S2.1 Stochastic behaviour of the Punisher alongside a single synthetic burdensome gene

In the main text's Figure 1C–D, we display the deterministically simulated the performance of the Punisher when our design is employed to penalise mutations of a single constitutive synthetic gene expressed by the cell (see Supplementary Note S1.5.1 for the constitutive gene expression ODEs and parameters). Here, we simulate the same scenario, but investigate the stochastic behaviour of our synthetic gene expression system using a hybrid tau-leap simulation algorithm [1]. In this approach, synthetic mRNA and protein levels are treated as stochastic variables but the cell's native gene expression is considered deterministically, because the host cell model's coarse-grained variables represent average dynamics of many species' concentrations, whose random fluctuation cancel each other out.

Namely, in Supplementary Figure S1 we show 50 stochastic trajectories for the host cell growth rates and the concentrations of the integrase and the CAT protein in the cell. The behaviour of the system is largely consistent with deterministic simulations, save for the fact that due to the stochasticity of gene expression, in some cases the Punisher becomes randomly switched on even before the burdensome gene is mutated. Another difference is the lower growth rate of the 'punished' mutant cells whose CAT gene copies have been excised by the integrase. This is because in our hybrid simulation the CAT gene DNA abundance $c_{cat}$ is a discrete integer variable that eventually becomes zero. In the deterministic simulations, however, $c_{cat}$ is a continuous variable governed by ODE (S20), which asymptotically decays towards zero but still remains somewhat positive, if small. Thus, deterministic simulations involve having non-zero antibiotic resistance protein levels in mutant cells, hence the higher cell growth rates in the main text's Figure 1C than in Supplementary Figure S1A. Given that in reality the number of gene copies per cell is indeed discrete and not continuous, we can expect the Punisher's real-life implementation to behave more similarly to our stochastic simulations, penalising mutant cells even more effectively than predicted by deterministic modelling.

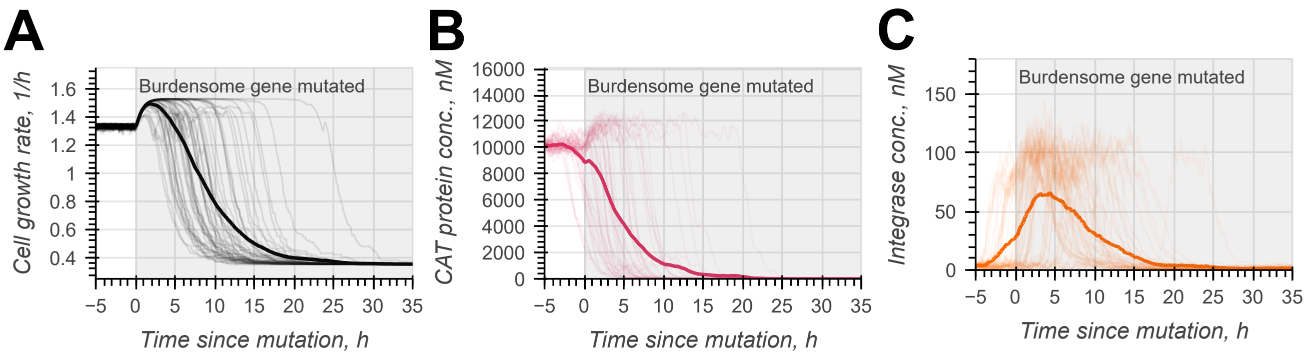

Supplementary Figure S1: Stochastic simulation of the Punisher's response to the mutation of a single constitutive burdensome gene expressed in the same host cell with it.

#### S2.2 Mutating one gene of two toggle switches present alongise the Punisher

In the main text's Figure 3B–C, we show how the Punisher penalises the mutation of both genes of one of the two synthetic toggle switches present in the cell together with it. In this supplementary note, we examine what happens if just one toggle switch gene becomes non-functional. As shown in Supplementary Figure S2A, mutating one toggle gene upregulates its counterpart in the same toggle switch, as its expression is no longer repressed. This can in fact **increase** the overall synthetic protein synthesis in the cell. Hence, while the Punisher does not become switched on due to the burden not falling but rather becoming greater, this is in any case unnecessary as the mutant cell grow slower than the original engineered cell (Supplementary Figure S2B).

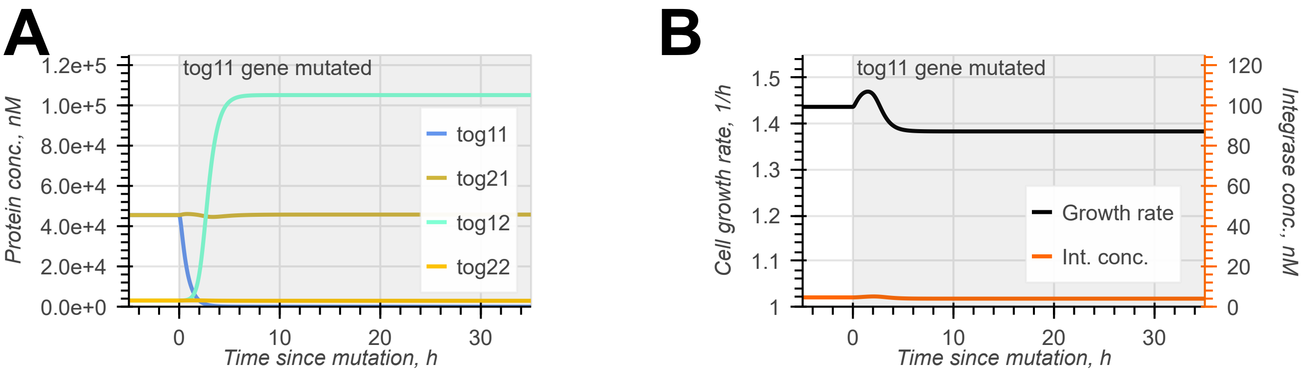

Supplementary Figure S2: Simulation of the Punisher's response to the mutation of a single toggle switch gene.

#### S2.3 Unwanted effects of essential co-expression on cells with synthetic toggle switch circuits

In the main text's Figure 4B–E, we compare the ability of different strategies to decrease the growth of cells hosting two synthetic toggle switch circuits, in which some toggle genes have been mutated. It was found that the co-expression, strategy in which the toggle genes are expressed from the same operon with an essential antibiotic resistance gene CAT, may sometimes result in less of a growth rate decrease (or possibly even boost the cell's growth) compared to that experienced by the cells with no synthetic circuitry besides the toggle switches themselves. This was especially true when the RBS sequences of the co-expressed CAT genes were weak. Thus, here we simulate such scenarios with the slow mRNA-ribosome association rates of $k^{+}_{cat_{yz}} = 0.24\ nM^{-1} \cdot h^{-1}$, i.e. $1/250$ of the maximum rate at the diffusion limit [6, 9, 10].

In Supplementary Figure S3 we see that, both with and without essential gene co-expression, mutating one toggle gene increases the overall burden experienced by the cell (the same can be said about the cell hosting our Punisher circuit alongside the toggles, shown in Supplementary Figure S2). However, in the case of co-expression, the CAT protein's concentration also increases. Whilst the mutant cells without any circuitry apart from the toggles still experience a greater burden than the original non-mutant cells and thus grow slightly slower, the increased CAT production in the co-expression case case confers mutants with a small growth advantage. Thus, essential gene co-expression with synthetic genes may unintentionally promote mutant cell growth, working directly against its design objective of slowing down mutant cell division.

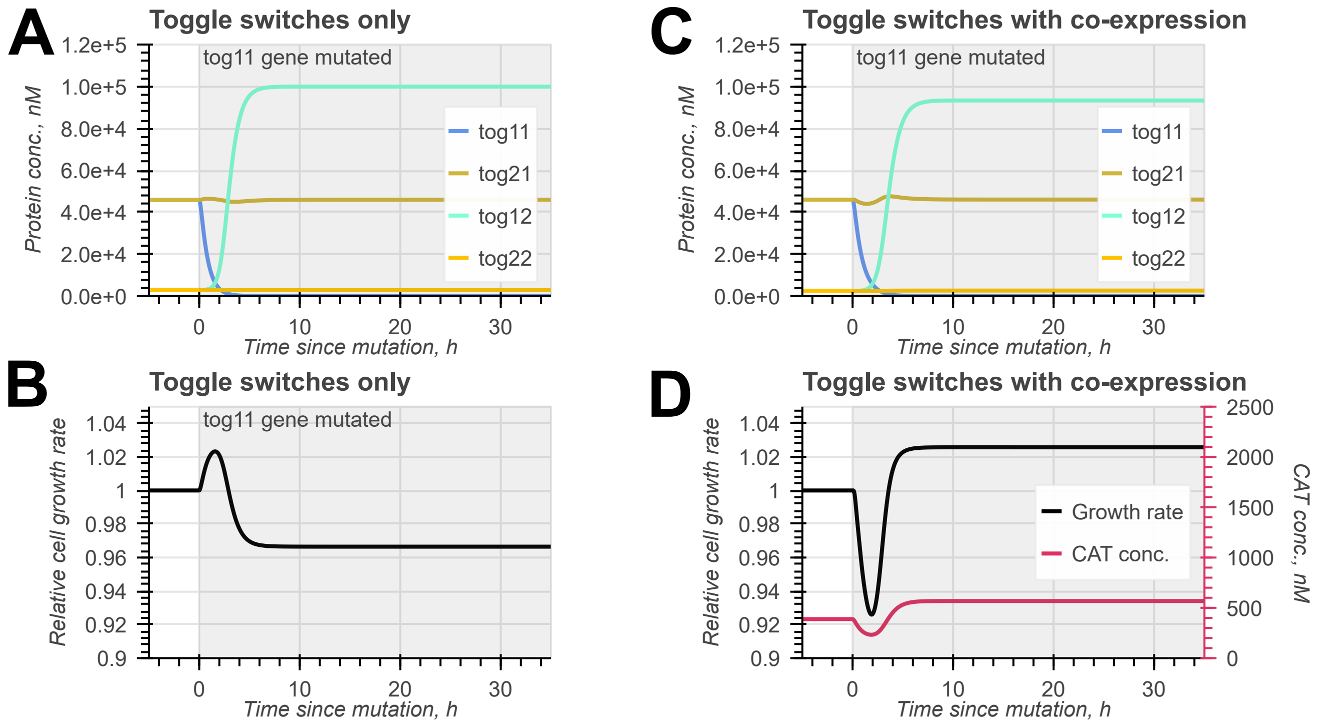

Supplementary Figure S3: Simulation of two synthetic toggle switches being hosted by the cell, with one gene of one toggle being mutated at $t = 0\ h$. Either **A–B)** no other synthetic circuitry is present in the cell or **C–D)** each of the four toggle switch genes is co-expressed with a CAT gene copy (see the main text's Figure 4A).

### S3 Switching threshold identification and tuning

This supplementary note outlines the analytical derivations made to determine the Punisher circuit's switching threshold – that is, the extent of competition for the cell's ribosomes (i.e. gene expression burden) at which our circuit transitions from expressing very little integrase to expressing a lot of it so as to excise an essential gene, penalising the mutated cell. To this end, we first state and motivate the simplifications made to our cell model as part of our analysis. Then, we introduce a lumped parameter which quantifies the burden experienced by the cell and treat it as a bifurcation parameter to show how the Punisher's possible steady state depend on resource competition. With the switching threshold identified, we determine the minimum increase in the rate of essential gene excision rate by the integrase associated with the Punisher becoming switched on. We also show it can be kept in place whilst the timescale of the Punisher is being tuned to suit a particular application.

#### S3.1 Simplifying assumptions

The analytical derivations that follow rely on two simplifying assumptions made about our cell model. First, we state that the total synthetic protein degradation flux is negligible compared to the total protein translation flux in the cell, i.e. $\Delta \ll \epsilon B$. This is not unreasonable, as in all the scenarios considered by us only the synthetic switch protein is degraded. Even if the entirety of the switch protein's synthesis flux was to be funnelled into the degradation flux (which it is not, since proteins are also removed by dilutio due to cell division), this would only constitute a small fraction of the overall protein synthesis. Indeed, the switch gene's maximum expression rate is $\approx$ 100 times smaller than that of the cell's metabolic gene class alone (compare $c_s\alpha_s$ in Supplementary Table S3 and $c_a\alpha_a$ in Supplementary Table S1), and metabolic proteins themselves never comprise more than a half of the cell's proteome. Hence, our first assumptions allows to approximate the cell growth rate and the resource competition denominator using Equations (S60) and (S61), where in Equation (S60) we use the fact that the abundance of actively translating ribosomes $B$ is included in the total ribosome abundance $R$, hence $\Delta \ll \epsilon R$.

$$D = \frac{K_D + h}{K_D} \cdot \left( 1 + \frac{\sum_{j \in \{a,r\} \cup X} m_j/k_j - \frac{K_D+h}{K_D} \cdot \frac{\overline{\phi}_q \Delta}{\epsilon R}}{1 - \overline{\phi}_q \left(1 - \frac{K_D+h}{K_D} \cdot \frac{\Delta}{\epsilon R}\right)} \right) \Leftrightarrow$$

$$\Leftrightarrow D \approx \frac{K_D + h}{K_D} \cdot \left(1 + \frac{\sum_{j\in\{a,r\}\cup X} m_j/k_j}{1-\overline{\phi}_q}\right) \tag{S60}$$

$$\lambda = \frac{\epsilon B}{M} - \frac{\Delta}{M} \approx \frac{\epsilon B}{M} = \frac{\epsilon \cdot R(\frac{K_D+h}{K_D} - \frac{1}{D})}{M} \Leftrightarrow$$

$$\Leftrightarrow \lambda \approx \frac{\epsilon R}{M} \cdot \frac{\frac{1}{1-\overline{\phi}_q} \sum_{j\in\{a,r\}\cup X} m_j/k_j}{\frac{K_D+h}{K_D} \cdot \left(1 + \frac{1}{1-\overline{\phi}_q} \sum_{j\in\{a,r\}\cup X} m_j/k_j\right)} \tag{S61}$$

The second simplification is that the steady-state values of the translation elongation rate $\epsilon$ and the ribosomal gene transcription regulation function $F_r$ are almost burden-independent. We also assume that this is the case, when the CAT gene is present and functional, for the fraction of ribosomes not bound by chloramphenicol

$$H = \frac{K_D}{K_D + h} \tag{S62}$$

Thus, for a given nutirent quality $\sigma$, extracellular chloramphenicol concentration $h_{ext}$ and set of *cat* gene expression parameters, the steady-state values $\overline{\epsilon}$, $\overline{F}_r$ and $\overline{H}$ can be found by running a simulation and thereafter treated as constant parameters. This also means that the effective mRNA-ribosome dissociation constants $\{k_j(\epsilon)\}\ \forall j \in \{a, r\} \cup X$ also become constant parameters $\{\overline{k}_j\}$.

We vindicate this second assumption by simulation. In order to provide a minimal working example, we only append the host cell model Equations (S1)–(S7) with the ODEs for the expression of CAT and a constitutive burdensome gene $b$ – that is, with Equations (S14) and (S18) and Equations (S41)–(S42) respectively. For a range of burdensome gene promoter strengths from $\alpha_b = 0$ (no synthetic gene expression except for CAT) to $\alpha_b = 2.5 \cdot 10^5$ (more synthetic gene expression than in any of the scenarios considered in this study), we retrieve the variables of interest's steady-state values by integrating the model ODEs over 50 $h$ of simulated time. Plotting the outcome of these simulations in Supplementary Figure S4, we see that for a wide range of culture medium nutrient qualities [6] even the highest burden does not affect the equilibrium values of $\epsilon$, $F_r$ and $H$ by more than 6%.

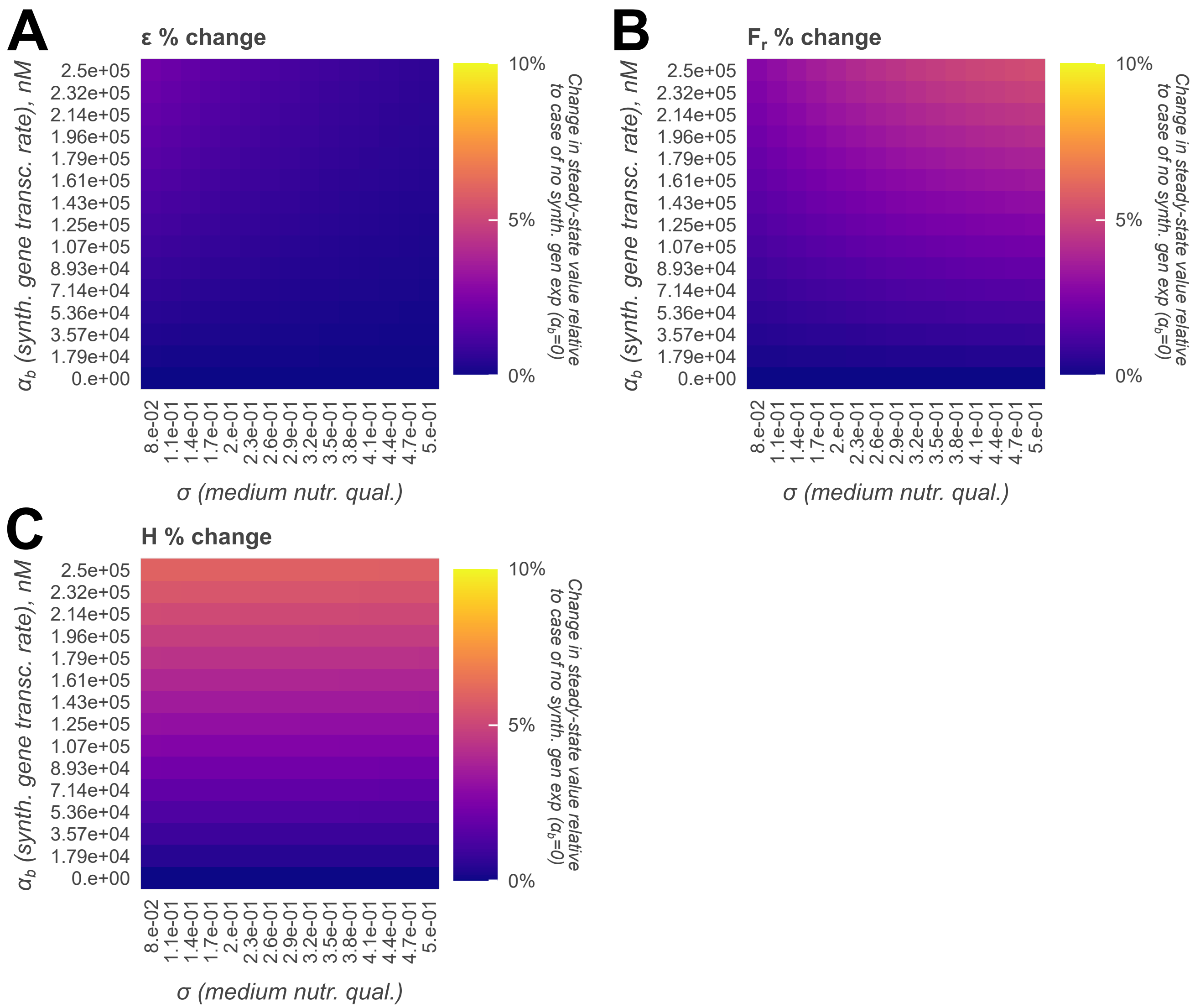

Supplementary Figure S4: Burden-induced changes in the cell's **A** translation elongation rate, **B** ribosome transcription regulation, **C** fraction of ribosomes free from chloramphenicol relative to the case of zero synthetic protein expression for different culture medium nutrient qualities (lowest quality $\sigma = 0.08$: M63 medium with glycelol as the carbon source; highest quality $\sigma = 0.5$: Rich Defined Medium with glucose as the carbon source [6]).

#### S3.2 Burden quantification

Leveraging the simplifications in Supplementary Note S3.1, let us now retrieve the circuit's steady state analytically. Equation S63 defines the equilibrium mRNA levels in terms of gene expression parameters, the steady-state cell growth rate $\overline{\lambda}$ and the steady-state value of gene $j$'s transcription regulation function $\overline{F}_j$.

$$\dot{m}_j = 0 \Leftrightarrow m_j = \frac{\overline{F}_j \alpha_j c_j \overline{\lambda}}{\overline{k}_j(\beta_j + \overline{\lambda})} \quad \forall j \in \{a, r\} \cup X \tag{S63}$$

Plugging Equations S63 and (S60)–(S61) into Equations S3, (S4) and (S9), we find that the steady-state concentrations of proteins which are not degraded by the Punisher's protease can be found as follows:

$$\dot{p}_j = 0 \Leftrightarrow \frac{\epsilon}{n_j} \cdot \frac{m_j/k_j}{D} - \overline{\lambda} p_j = 0 \Leftrightarrow$$

$$\Leftrightarrow p_j = \frac{M(1-\overline{\phi}_q)}{n_j} \cdot \frac{\dfrac{\overline{F}_j \alpha_j c_j \overline{\lambda}}{\overline{k}_j(\beta_j + \overline{\lambda})}}{\displaystyle\sum_{l \in \{a,r\} \cup X} \dfrac{\overline{F}_l \alpha_l c_l \overline{\lambda}}{\overline{k}_l(\beta_l + \overline{\lambda})}} \quad \forall j \in \{a,r\} \cup \{x_l \in X : \delta_{x_l} = 0\} \tag{S64}$$

Now, we note that mRNA degradation rates are roughly identical across all genes: indeed, none of the circuits considered whilst analysing the Punisher's performance involve mRNA removal through anything but overall RNA degradation (for instance, there is no RNA-based antithetic integral feedback). Hence, Equation (S64) simplifies to:

$$p_j = \frac{M(1-\overline{\phi}_q)}{n_j} \cdot \frac{\overline{F}_j \alpha_j c_j / \overline{k}_j}{\displaystyle\sum_{y \in \{a,r\} \cup X} \overline{F}_y \alpha_y c_y / \overline{k}_y} \quad \forall j \in \{a,r\} \cup \{x_l \in X : \delta_{x_l} = 0\} \tag{S65}$$

Equation (S65) reveals that the steady-state concentration of a given protein depends on the total resource competition from all the genes in the cell, which is reflected by the denominator. It is then natural to state that

$$\xi_j = \frac{\overline{F}_j c_j \alpha_j}{\overline{k}_j} \tag{S66}$$

quantifies the burden imposed by the gene $j \in \{a,r\} \cup X$ on the cell. Equation (S65) can thus be rewritten as Equation (S67), which shows that the steady-state concentration of a given gene's protein is defined by the ratio of the competition for ribosomes that it imposes on the cell to the total resource competition exerted by all native and synthetic genes in the cell. This denominator can therefore be viewed as the host cell's overall gene expression burden.

$$p_j = \frac{M(1-\overline{\phi}_q)}{n_j} \cdot \frac{\xi_j}{\displaystyle\sum_{l \in \{a,r\} \cup X} \xi_l} \quad \forall j \in \{a,r\} \cup \{x_l \in X : \delta_{x_l} = 0\} \tag{S67}$$

#### S3.3 Circuit steady state retrieval

Having expressed the gene expression burden experience by the cell as a parameter, let us now investigate how it determines the steady states of the Punisher circuit, and specifically the equilibrium concetration of the switch protein. Since it is degraded by the synthetic protease, the steady-state $p_s$ value is not given straightforwardly by Equation (S67). However, Equation (S67) does yield the concentration of the protease itself and of the cell's ribosomes, which we can then plug into ODE (S15) to get:

$$\dot{p}_j = 0 \Leftrightarrow \frac{\epsilon}{n_j} \cdot \frac{m_j/k_j}{D} \cdot R - (\delta_s p_{prot} + \overline{\lambda})p_j = 0 \Leftrightarrow$$

$$\Leftrightarrow p_s = \frac{M(1-\overline{\phi}_q)}{n_s} \cdot \frac{\xi_s}{\sum\limits_{l\in\{a,r\}\cup X} \xi_l} \cdot \left(1 + \frac{M\delta_s}{\overline{H}\overline{\epsilon}} \cdot \frac{n_r \xi_{prot}}{n_{prot}\xi_r} \cdot \frac{1 + \frac{1}{1-\overline{\phi}_q}\sum_{j\in\{a,r\}\cup X} m_j/k_j}{\frac{1}{1-\overline{\phi}_q}\sum_{j\in\{a,r\}\cup X} m_j/k_j}\right)^{-1} \tag{S68}$$

Notably, one of the denominator's terms includes the quotient

$$\frac{1 + \frac{1}{1-\overline{\phi}_q}\sum_{j\in\{a,r\}\cup X} m_j/k_j}{\frac{1}{1-\overline{\phi}_q}\sum_{j\in\{a,r\}\cup X} m_j/k_j} = 1 + \frac{1}{\frac{1}{1-\overline{\phi}_q}\sum_{j\in\{a,r\}\cup X} m_j/k_j} \tag{S69}$$

which involves the reciprocal of the sum of all mRNA concentrations scaled by the apparent mRNA-ribosome dissociation constants. For the host cell's native genes, these concentrations are on the order of magnitude of $10^4$ - $10^5$ $nM$, whereas the dissociation constants have the order of magnitude of $10^0$ - $10^1$ $nM$ (see Supplementary Tables S1–S2 and [1]). Hence, the denominator in Equation (S69) is a sum of very large ${}^{m_a}/_{k_a}$ and ${}^{m_r}/_{k_r}$ and additional non-negative terms for synthetic mRNAs, scaled by the coefficient $\frac{1}{1-\overline{\phi}_q} = \frac{1}{0.41} > 1$. It is thus reasonable to state that $\frac{1}{1-\overline{\phi}_q}\sum_{j\in\{a,r\}\cup X} m_j/k_j \gg 1$, which reduces the quotient in question to just 1. An estimate for the steady-state $p_s$ value is thus given by Equation (S70).

$$p_s \approx \frac{M(1-\overline{\phi}_q)}{n_s} \cdot \frac{\xi_s}{\sum\limits_{y\in\{a,r\}\cup X} \xi_y} \cdot (1+\chi)^{-1} \tag{S70}$$

$$\text{where } \chi = \frac{M\delta_s}{\overline{H}\overline{\epsilon}} \cdot \frac{n_r \xi_{prot}}{n_{prot}\xi_r} \tag{S71}$$

However, this equation is still not sufficient to find the equilibrium value of $p_s$ from the gene circuit's design parameters, as according to Equation (S66) the switch protein expression burden $\xi_s$

depends on the switch gene's steady-state transcription regulation function $\overline{F}_s$, which has not been determined yet. Since the integrase is co-expressed with the switch gene from the same operon, a similar issue arises when we attempt retrieving the integrase expression burden $\xi_i$. To work around this problem, we reformulate our question. Instead of trying to find the equilibrium value of $p_s$, we ask: provided that $p_s$ **is** the equilibrium switch protein concentration, what steady-state value of the transcription regulation function $\overline{F}_s$ is **required** to achieve it?

Given that $0 \leq \overline{F}_s \equiv \overline{F}_i \leq 1$ (see Equation S19), the values $\xi_s$ and $\xi_i$ can be expressed as

$$\xi_s = \overline{F}_s \xi_s^{max}, \; \xi_i = \overline{F}_s \xi_i^{max} \tag{S72}$$

where

$$\xi_s^{max} = \frac{c_s \alpha_s}{\overline{k}_s} \text{ and } \xi_i^{max} = \frac{c_i \alpha_i}{\overline{k}_i} \equiv \frac{c_s \alpha_s}{\overline{k}_i} \tag{S73}$$

are constant lumped parameters which can indeed be found straightforwardly from the Punisher circuit's design parameters. This allows to rewrite Equation (S70) into Equation (S74), where $\Xi$ is the gene expression burden due to all genes except for the Punisher's switch and integrase genes.

$$p_s = \frac{M(1-\overline{\phi}_q)}{n_s} \cdot \frac{\overline{F}_s \xi_s^{max}}{\Xi + \overline{F}_s(\xi_s^{max} + \xi_i^{max})} \cdot (1+\chi)^{-1} \tag{S74}$$

$$\text{where} \quad \Xi = \sum_{j \in \{a,r\} \cup X \setminus \{s,i\}} \xi_j \tag{S75}$$

Solving Equation (S74) for $\overline{F}_s$, we finally obtain Equation (S76), which allows to calculate the transcription regulation function value that is required to achieve a steady-state switch protein abundance $p_s$ under a given gene expression burden $\Xi$.

$$\overline{F}_s^{req}(p_s, \Xi) = p_s(1+\chi) \cdot \frac{\Xi}{\xi_s^{max} + \xi_i^{max}} \cdot \left( \frac{M(1-\overline{\phi}_q)}{n_s} \cdot \frac{\xi_s^{max}}{\xi_s^{max} + \xi_i^{max}} - p_s(1+\chi) \right)^{-1} \tag{S76}$$

However, the value of $F_s$ is in reality a function of $p_s$, given by the Hill function in Equation (S19). We denote it as the 'real' transcription regulation function value $\overline{F}_s^{real}(p_s)$. The sufficient and necessary condition for having a given equilibrium switch protein level $p_s$ is therefore outlined in Equation (S77), which clearly demonstrates that the nature of the system's equilibria depends on the burden $\Xi$.

$$\overline{F}_s^{req}(p_s,\Xi) = \overline{F}_s^{real}(p_s) = F_{s_b} + (1-F_{s_b}) \cdot \frac{(Ip_s)^{\eta_s}}{(Ip_s)^{\eta_s} + K_s^{\eta_s}} \tag{S77}$$

Plotting the real and required values of $F_s$ in the main text's Figure 2A therefore allows to find the system's equilibria, as well as their stability: the switch protein's abundance decreases if the real value of $F_s$ for this concentration $p_s$ is lower than the one required to maintain it as a steady-state value, and vice versa.

#### S3.4 Switching threshold identification

We want the Punisher to 'switch on' – that is, transition from an equilibrium with low switch protein (and integrase) expression to a high-expression equilibrium – when a synthetic gene mutation occurs and reduces burden below a threshold value. From our definition of $\Xi$ in Equation (S75), we can see that such mutation of a synthetic gene $x_l$ stands for subtracting the mutated gene's contribution from the total burden parameter, i.e.

$$\text{Gene } x_l \text{ mutated } \Leftrightarrow \Xi \text{ reduced from } \Xi_0 \text{ to } (\Xi_0 - \xi_{x_l}) \tag{S78}$$

Knowing Equation (S77), it is thus easy to understand that a 'switching' change in the identity of the Punisher's equilibria is underlain by a **saddle node bifurcation** caused by the change in $\Xi$. Indeed, when burden is decreased, the system goes from having an unstable and two stable equilibria – one at a high $p_s$ value, another at a low one (which is the one the Punisher is originally in) – to having one equilibrium at a high value of $p_s$ and a saddle node to its left. This saddle node then immediately vanishes, leaving only the high-expression stable steady state, when the burden is further reduced.

The saddle node stands for the point where the $\overline{F}_s^{req}(\Xi,p_s)$ curve touches the $\overline{F}_s^{real}$ curve from below (see the main text's Figure 2A). Mathematically, this point is defined as the solution to the following problem:

$$\overline{F}_s^{req}(p_s, \Xi) = \overline{F}_s^{real}(p_s) = F \tag{S79}$$

$$\frac{d\overline{F}_s^{real}}{dp_s} = \frac{d\overline{F}_s^{req}}{dp_s} \tag{S80}$$

$$\frac{d^2\overline{F}_s^{real}}{dp_s^2} > 0 \tag{S81}$$

Let us reformulate these conditions in terms of $F$, the common value of the required and real transcription regulation functions at the saddle point. By substituting $F = \overline{F}_s^{real}$ into Equation (S19), we can find $p_s$ as a function of $F$ according to Equation (S82).

$$p_s(F) = \frac{K_s}{I} \cdot \left(\frac{F - F_{s_b}}{1 - F}\right)^{1/\eta_s} \tag{S82}$$

which then allows to solve Equation (S76) for $F = \overline{F}_s^{req}$ to define $\Xi$ as

$$\Xi(F) = F \cdot \frac{\xi_s^{max} + \xi_i^{max}}{(1+\chi)p_s(F)} \cdot \left(\frac{M(1-\overline{\phi}_q)}{n_s} \cdot \frac{\xi_s^{max}}{\xi_s^{max} + \xi_i^{max}} - (1+\chi)p_s(F)\right) \tag{S83}$$

Now, let us observe that by defimition protein concentrations cannot be negative, which bounds $p_s$ from below. On the other hand, differentiating the expression for $\overline{F}_s^{req}$ (Equation (S19)) twice and substituting it into Condition (S81) bounds $p_s$ from above. Together, these considerations mean that $0 \le p_s \le \frac{K}{I} \cdot \left(\frac{\eta_s - 1}{\eta_s + 1}\right)^{1/\eta_s}$. Since $p_s(F)$ increases monotonically with $F$, this is equivalent to having the bounds:

$$F_{s_b} = \overline{F}_s^{req}(p_s = 0) \le F \le \overline{F}_s^{req}\left(p_s = \frac{K}{I} \cdot \left(\frac{\eta_s - 1}{\eta_s + 1}\right)^{1/\eta_s}\right) \tag{S84}$$

Differentiating $\overline{F}_s^{req}$ and $\overline{F}_s^{real}$ by $p_s$ and plugging $p_s(F)$ and $\Xi(F)$ into the obtained expressions, we can reformulate the problem of finding the desired saddle point as a one-dimensional root-finding problem shown below. Having found the saddle-node value $\hat{F}_s$, the corresponding switch protein concentration $\hat{p}_s$ and gene expression burden $\hat{\Xi}$ can be found according to Equations (S83) and (S82).

$$\textbf{Solve}\quad \eta_s(1-F_{s_b})\cdot\frac{\left(\frac{K_s}{I}\right)^{\eta_s}\cdot(p_s(F))^{\eta_s-1}}{\left(\left(\frac{K_s}{I}\right)^{\eta_s}+(p_s(F))^{\eta_s}\right)^2}=$$

$$=\frac{\Xi(F)\xi_s^{max}M(1-\overline{\phi}_q)}{n_s}\cdot\left(p_s(F)\cdot(\xi_s^{max}+\xi_i^{max})-\frac{\xi_s^{max}M(1-\overline{\phi}_q)}{n_s}\right)^{-2} \tag{S85}$$

**for**

$$F_{s_b}\leq F\leq \overline{F}_s^{req}\left(\frac{K}{I}\cdot\left(\frac{\eta_s-1}{\eta_s+1}\right)^{1/\eta_s}\right) \tag{S86}$$

#### S3.5 Minimum change in integrase activity

The Punisher's integrase is expressed, albeit at low levels, even when our circuit is not activated. Hence, even unmutated cells still experience a small rate of 'off-target' essential gene excision. However, despite this action non-mutant cells may still have a growth advantage over mutants, provided that the rate of essential gene excision once the Punisher actually **has** been activated by burden reduction must be significantly increased compared to the off-target rate. A minimum increase in the integrase's expression upon the Punisher being switched on can be found by looking at the saddle-node bifurcation point as shown in Supplementary Note S3.4. Activation of the punisher by synthetic gene mutations will therefore increase the integrase's expression **at least** by this retrieved minimum value, from which we can calculate the overall increase in the integrase's DNA-cutting activity.

An intuition why this is the case is provided by Supplementary Figure S5. Before the Punisher is switched on, it experiences a burden $\Xi > \hat{\Xi}$ that is above the bifurcation threshold, which gives rise to a low-expression equilibrium transcription regulation function value $F_s^{low}$. This is smaller than that at the saddle bifurcation point ($F_s^{low} < \hat{F}_s$). After the Punisher is switched on, the burden is below the threshold ($\Xi < \hat{\Xi}$), which gives rise to a single high-expression equilibrium at $F_s^{high}$. This transcription regulation function value lies above the second, non-saddle equilibrium $\hat{F}'_s$ for $\Xi = \hat{\Xi}$. Hence, the change in the transcription regulation function of the switch and integrase genes that accompanies the Punisher's switching is

$$F_s^{high}-F_s^{low} > \hat{F}'_s-\hat{F}_s \tag{S87}$$

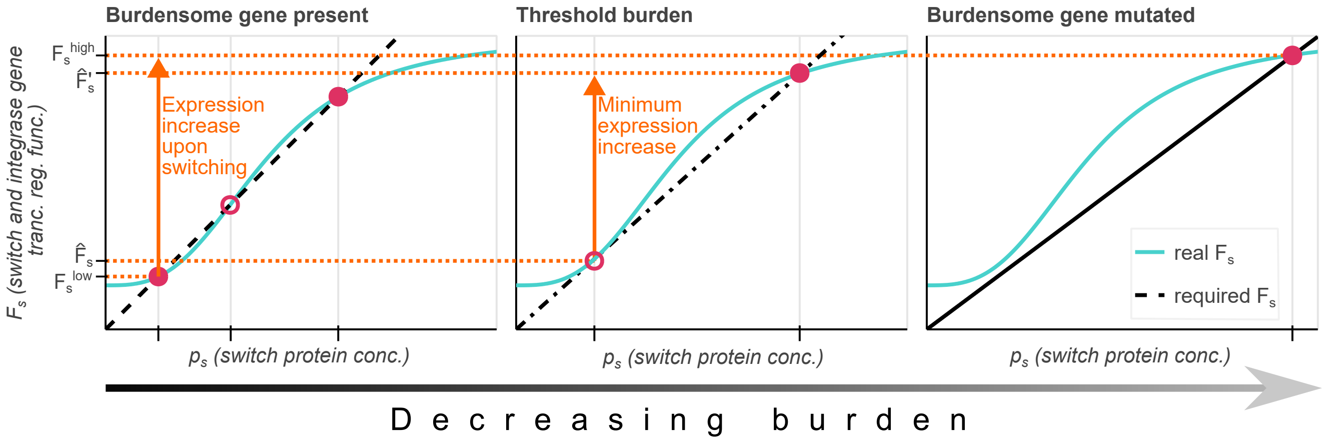

Supplementary Figure S5: When the Punisher is switched on by burdensome gene mutation, the integrase (and switch) genes' transcription regulation function increases from $F_s^{low}$ to $F_s^{low}$. Since the equilibria $\hat{F}_s$ and $\hat{F}'_s$ at the bifurcation point lie between these two values, the difference between them provides a minimum for this expression increase.

Thus, Equation (S87) introduces a lower bound for the change in the integrase gene's transcription regulation function upon the Punisher's activation. Using Equation (S67), for a given $F_s$ value the integrase's steady-state concentration can be found according to Equation (S88).

$$p_i(F_s, \Xi) = \frac{M(1-\overline{\phi}_q)}{n_i} \cdot \frac{F_s \xi_i^{max}}{\Xi + F_s(\xi_s^{max} + \xi_i^{max})} \tag{S88}$$

The integrase's activity in the cell can be calculated according to the expression in Equation (S89) [11], which increases monotonically as the integrase's abundance rises.

$$\frac{p_i^4}{p_i^4 + K_{bI}^4} \tag{S89}$$

Thus, if $\hat{F}'$ and $\hat{F}$ are known, the minimum fold-change in the integrase's activity is found as:

$$\left(\frac{\left(p_i(\hat{F}', \hat{\Xi})\right)^4}{\left(p_i(\hat{F}', \hat{\Xi})\right)^4 + K_{bI}^4}\right) \div \left(\frac{\left(p_i(\hat{F}, \hat{\Xi})\right)^4}{\left(p_i(\hat{F}, \hat{\Xi})\right)^4 + K_{bI}^4}\right) \tag{S90}$$

The retrieval of the saddle-node transcription regulation function value $\hat{F}$ and the corresponding threshold burden value $\hat{\Xi}$ has been discussed in Supplementary Note S3.4. Therefore, to calculate the desired minimum activity change, we only need to find the second, higher-expression, equilibrium $\hat{F}'$ achieved for the same burden.

We do this in two steps. First, we find the **switch** protein's concentration $p_s$ corresponding to this equilibrium. As mentioned in Supplementary Note S3.3, this is simply equivalent to solving Equation (S77). Since this higher-expression equilibrium is situated above the saddle node, this problem's domain is defined as the interval between the saddle-node equilibrium $p_s$ value and the maximum possible value of $p_s$, achieved when the gene's promoter is fully transcribed (i.e. $F_s = 1$). We formulate this one-dimensional root-finding problem below.

$$\textbf{Solve} \;\; \overline{F}_s^{req}(p_s, \hat{\Xi}) = \overline{F}_s^{real}(p_s) \tag{S91}$$

$$\textbf{for}$$

$$\frac{K_s}{I} \cdot \left( \frac{\hat{F} - F_{s_b}}{1 - \hat{F}} \right)^{1/\eta_s} \leq p_s \leq \frac{M(1 - \overline{\phi}_q)}{n_s} \cdot \frac{1 \cdot \xi_s^{max}}{\hat{\Xi} + 1 \cdot (\xi_s^{max} + \xi_i^{max})} \cdot (1 + \chi)^{-1} \tag{S92}$$

Second, we use Equation (S19) to find $\hat{F}'$ from the retrieved solution $\hat{p}'_s$ as:

$$\hat{F}' = \overline{F}_s^{real}(\hat{p}'_s) = F_{s_b} + (1 - F_{s_b}) \cdot \frac{(I\hat{p}'_s)^{\eta_s}}{(I\hat{p}'_s)^{\eta_s} + K_s^{\eta_s}} \tag{S93}$$

#### S3.6 Switching Timescale Adjustment

Supplementary Notes S3.3–S3.4 provide instructions for determining the Punisher's switching threshold value of burden for a given set of circuit and environmental parameters. As shown in the main text's Figure 2, this allows to establish the trends in how this threshold changes as different parameters are varied. In particular, one can alter the chemical induction of the switch gene's autoactivation (captured by the parameter $I$) to set a switching threshold that is appropriate for a given set of culture conditions and synthetic genes whose mutations are to be penalised. Notably, changing $I$ is not a genetic intervention, which means that adapting it for new applications is fast and cheap.

However, besides determining **whether** the Punisher is switched on by a certain fall in burden, it may also be important to adjust **how fast** this occurs. For instance, we may expect that some distrubances or oscillation may transiently reduce gene expression burden below the threshold for a limited time without an underlying synthetic gene mutation. In this case, we would like to make sure that by the time this transient perturbation subsides and the original high burden is restored, the Punisher will not have left its low-expression equilibrium's basin of attraction and will converge back to its 'off' state, rejecting the disturbance.

One way to change the speed of switching would be to readjust the threshold burden value itself, decreasing it but still keeping it above the burden experienced by mutant cells (Supplementary Figure S6A–B). However, this may prove challenging if the application in question has a narrow range of suitable threshold values. Hence, this supplementary note proposes a means of changing the timescale of the Punisher's activation whilst keeping the switching threshold burden value $\hat{\Xi}$ unchanged.

The timescale of a biological system's dynamics is primarily determined by the rate of the removal of molecules [13] – which in our case would be the total rate of dilution and degradation of the switch protein. Whilst dilution happens due to cell division and thus cannot be controlled directly, the switch protein degradation rate can be easily adjusted either by changing the abundance of the synthetic protease in the cell (e.g. by having its gene be expressed) or by altering the protease degradation tag in the switch protein's coding sequence. Whilst the latter intervention by definition involves DNA editing, only a few codons need to be altered at a time – for instance, in [7] protein degradation rates could be varied up to $\approx$ 20-fold by using different amino acids in just six positions. Such small-scale DNA edits can be done using cheap and efficient techniques like CRISPR as opposed to cumbersome assembly of large DNA fragments from scratch [14, 15].

Now, suppose that we have a Punisher circuit with a switching threshold already set to be appropriate for a particular application. In order to change the timescale of its activation, we alter the protease's expression or the switch protein's affinity to it by editing the tag sequence. How can we now ensure that this does not affect the circuit's switching threshold burden value $\hat{\Xi}$?

From Supplementary Note S3.3 that the circuit's steady states and bifurcations are determined by solving Equation (S94) in terms of the switch protein's concentration $p_s$ and the burden parameter $\Xi$.

$$\overline{F}_s^{req}(p_s,\Xi)=\overline{F}_s^{real}(p_s)\Leftrightarrow$$
$$\Leftrightarrow\frac{p_s(1+\chi)\Xi}{\xi_s^{max}+\xi_i^{max}}\cdot\left(\frac{M(1-\overline{\phi}_q)}{n_s}\cdot\frac{\xi_s^{max}}{\xi_s^{max}+\xi_i^{max}}-p_s(1+\chi)\right)^{-1}=F_{s_b}+(1-F_{s_b})\cdot\frac{(Ip_s)^{\eta_s}}{(Ip_s)^{\eta_s}+K_s^{\eta_s}} \tag{S94}$$

Examining this equation, we observe that on the right-hand side, $p_s$ is always scaled by the chemical induction parameter $\Xi$, whereas on the left-hand side it is always scaled by $(1+\chi)$, where

$$\chi = \frac{M\delta_s}{\overline{H}\overline{\epsilon}} \cdot \frac{n_r \xi_{prot}}{n_{prot}\xi_r} \tag{S95}$$

is the coefficient capturing the effects of the switch protein's degradation by the protease. Notably, Equation (S95) includes both the degradation tag sequence strength in the form of the degradation rate parameter $\delta_s$ and the extent of protease gene expression in the form of the translational burden $\xi_{prot}$ defined as a combination of the protease gene's concentration and promoter and RBS strengths.

Thus, if the original fine-tuned Punisher had parameters $I_0$ and $\chi_0$ but our adjustment of $\delta_s$ or the protease gene expression parameters has given rise to a new $\chi = \chi_1$, the solution to Equation (S94) in terms of $\Xi$ (and thus the Punisher's switching threshold) will remain the same as in the original case as long as the new chemical induction parameter $I_1$ satisfies Equation (S96).

$$\frac{I_1}{I_0} = \frac{1+\chi_1}{1+\chi_0} \tag{S96}$$

To see why this is the case, recall that graphically solutions to Equation (S94) can be understood as intersections between the curves representing the transcription regulation functions $\overline{F}_{req}$ and $\overline{F}_{real}$ plotted against the corresponding values of $p_s$. There, the burden $\Xi$ determines the slope of the $y = \overline{F}_{req}(p_s = x)$ curve (see Figure S5 and the main text's Figure 2A). Scaling $p_s$ in both functions by the same factor $\frac{I_1}{I_0} = \frac{1+\chi_1}{1+\chi_0}$ is therefore equivalent to applying a coordinate transform to the horizontal axis for both of the curves. The slope of $\overline{F}_{req}$ that makes the curves intersect exactly twice (which produces the saddle-node bifrucation, which corresponds to the Punisher being switched on) will therefore be the same before and after the coordinate transform. Hence, Equation (S96) ensures that the switching threshold value of burden $\hat{X}i$ remains the same.

An example of adjusting the Punisher's timescale of switching is displayed in Figure S6C–H. Here, we do it for the same scenario as the one considered in the main text's Section 3 and described in detail in Supplementary Note S1.5.2 – that is, the Punisher being deployed to penalise mutations of synthetic two toggle switch circuits hosted by the cell. However, this time we set a higher switching threshold than before (although this threshold is still below the burden of expessing all toggle switch genes). Hence, when two toggle switches are flipped by an inducer concentration pulse, the transient reduction in burden for a limited time is sufficient to make the

Punisher leave its low-expression equilibrium's basin of attraction. Thus, by the time the high burden is restored, the punished has already been switched on irreversibly, which causes it to wrongly penalise a genetically intact cell (Supplementary Figure S6C–D).

We therefore adjust the switch protein's degradation tag to one with a weaker affinity for the protease, which yields a new switch protein degradation rate $\delta_s = 0.001934\ nM^{-1} \cdot h^{-1}$ instead of $\delta_s = 0.01836\ nM^{-1} \cdot h^{-1}$ [7]. In line with Equation (S96), we change the chemical induction level from $I = 0.88$ to $I = 0.192$. As shown in Supplementary Figure S6E–F, this prevents the 'false positive' irreversible switching on of the Punisher in response to transient fluctuations in burden. More generally, by showing the switching timescale's dependence on the parameters $\delta_s$ and $I$ in Supplementary Figure S6G, we show how the same switching threshold is conserved along the entire white line defined by Equation (S96). Meanwhile, Supplementary Figure S6H shows that moving along this line allows to significantly alter the time taken by the Punisher to become irreversibly switched on by a reduction in burden.

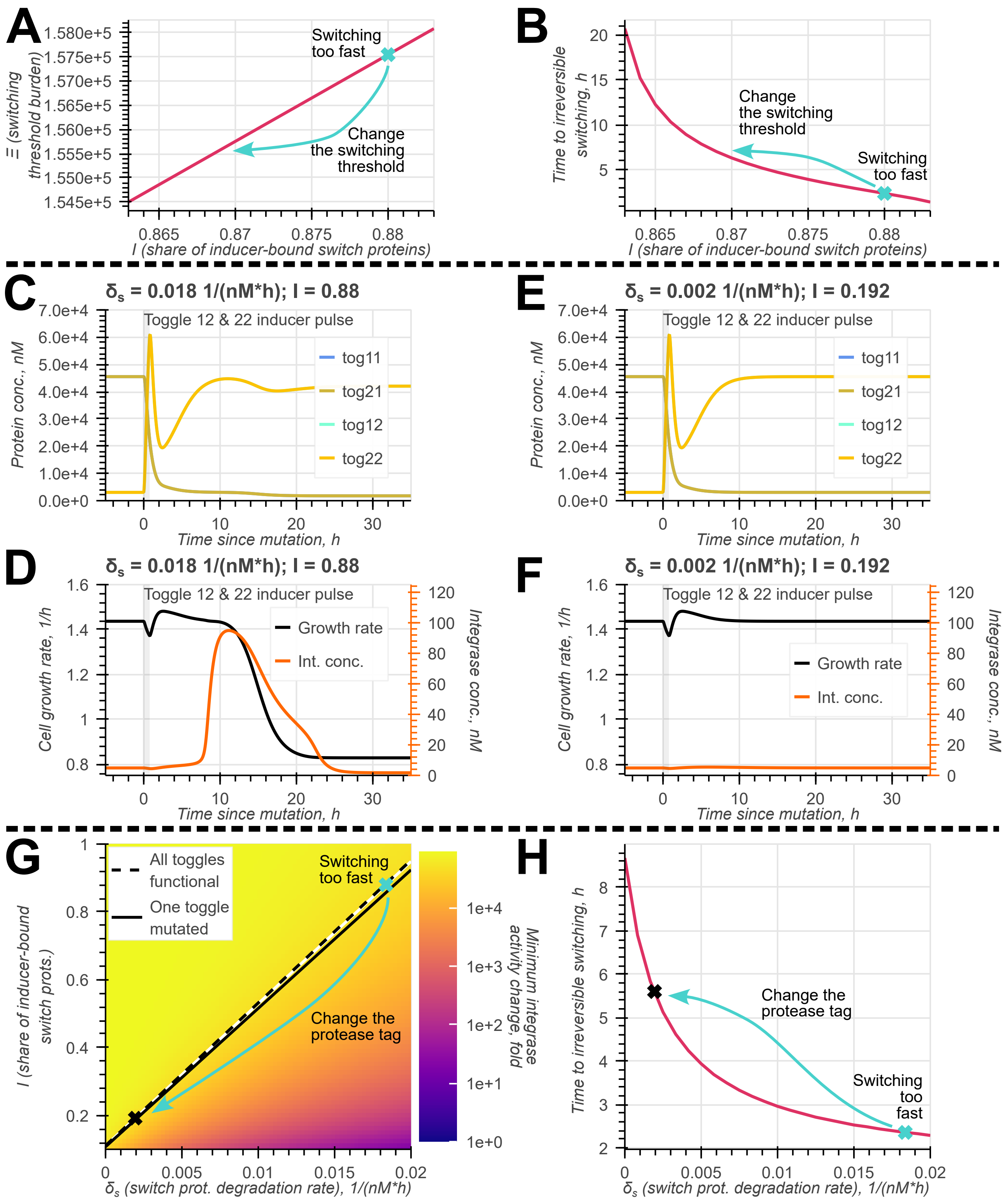

A
Switching too fast
Change the switching threshold
Ξ (switching threshold burden)
I (share of inducer-bound switch proteins)
B
Time to irreversible switching, h
Change the switching threshold
Switching too fast
I (share of inducer-bound switch proteins)
C
δs = 0.018 1/(nM*h); I = 0.88
Toggle 12 & 22 inducer pulse
tog11
tog21
tog12
tog22
Protein conc., nM
Time since mutation, h
E
δs = 0.002 1/(nM*h); I = 0.192
Toggle 12 & 22 inducer pulse
Protein conc., nM
Time since mutation, h
D
δs = 0.018 1/(nM*h); I = 0.88
Toggle 12 & 22 inducer pulse
Growth rate
Int. conc.
Cell growth rate, 1/h
Integrase conc., nM
Time since mutation, h
F
δs = 0.002 1/(nM*h); I = 0.192
Toggle 12 & 22 inducer pulse
Growth rate
Int. conc.
Cell growth rate, 1/h
Integrase conc., nM
Time since mutation, h
G
All toggles functional
One toggle mutated
Switching too fast
Change the protease tag
Minimum integrase activity change, fold
I (share of inducer-bound switch prots.)
δs (switch prot. degradation rate), 1/(nM*h)
H
Time to irreversible switching, h
Change the protease tag
Switching too fast
δs (switch prot. degradation rate), 1/(nM*h)

Supplementary Figure S6: The effect and tuning of the Punisher's switching timescale. **A–B** Shifting the Punisher's switching threshold burden value $\hat{\Xi}$ by varying the chemical induction of the switch gene $I$ allows to adjust the time of the Punisher's irreversible switching (i.e. time until it gets out of the off-state's low-expression equilibrium's basin of attraction). The lower is the switching threshold (as long as it is still above the burden experienced by mutated cells), the longer the Punisher takes to become switched on irreversibly. **C–D** A fast switching timescale of the punihser means that it can get triggered by a temporary reduction in burden associated with the flipping of the synthetic toggle switch circuits. **E–F** Reducing the rate of the switch protein's degradation by the protease and readjusting the chemical inducer concentration according to Equation (S96) prevents triggering of the Punisher in response to transient fluctuations in burden. **G** The effect of the chemical induction and the switch protein degradation rate on the Punisher's switching threshold and the minimum fold-increase in the integrase's activity upon the Punisher being switched on (see Supplementary Note S3.5). The blue cross stands for the fast-switching parameter combination in (C–D) and the black cross stands for the adjusted parameters in (E–F). All parameter combinations with the same switching threshold burden value as in (C–F) are given by the white line, defined according to Equation (S96). **H** The time of the Punisher's irreversible switching (i.e. time until it gets out of the off-state's low-expression equilibrium's basin of attraction) for different switch protein degradation rates. For each $\delta_s$ value, the value of $I$ was adjusted according to Equation (S96) so as to maintain the same switching threshold. This is equivalent to moving along the white line in E. The blue cross stands for $\delta_s$ and irreversible switching time in (C–D) and the black cross stands for these values in (E–F).

### S4 Cell population model

In the main text's Section 4, we simulate the Punisher's effect on the genetic stability of a population of cells grown in a bioreactor, which allows to evaluate if our design fulfils its intended task of mitigating mutation spread in engineered cell cultures. This supplementary note explains the population model was used to perform the above simulations, as well as outlines how it was parameterised based on simulations of our single-cell coarse-grained resource-aware model described in Supplementary Note S1.

#### S4.1 Population state vector

Our population model is defined for the simplest use case for the Punisher (described in the main text's Section 2.1 and Supplementary Note S1.5.1), where the only synthetic gene present in the host cell apart from the Punisher itself is a constitutive burdensome gene. In order to simplify the consideration of gene mutations, we assume that only a single copy of each Punisher gene is present in the cell (instead of having copy numbers of 10 $nM$ as in our single-cell simulations). To maintain the same total gene transcription rates as before despite this change, the promoter strengths $\alpha_s$, $\alpha_i$, $\alpha_{prot}$ and $\alpha_{cat}$ were increased tenfold compared to their values in Supplementary Table S3.

As discussed in the main text's Section 4, we divide the cell population into several subpopulations based on the state of the Punisher circuit in the cell and which of the cell's synthetic genes, if any, have been rendered non-functional – i.e. unable to express the encoded protein – by mutation. The number of cells in each subpopulation is treated as a deterministic variable. These cell counts $\{x_j\}$ are gathered into the vector $\mathbf{x} = (x_1\ x_2\ \ldots x_N)^\intercal$, whose dynamics are simulated by ODE integration. Note that for the sake of simplicity, we leverage the fact that the switch and the integrase are co-expressed from the same operon and assume that either both of them are functional or both of them are not expressed. This allows us to reduce the number of subpopulations considered by our model.

For convenience, instead of numbers standing for positions in the $\boldsymbol{x}$ vector, it can be useful to index the state vector's entries based on the cell subpopulations they represent. An index $j$ therefore has the form:

$$j = G_1G_2G_3G_4 : G_5$$

where

$$G_1 = \begin{cases} B, \text{ if the synthetic burdensome gene is functional} \\ B', \text{ if the burdensome gene has been mutated} \end{cases}$$

$$G_2 = \begin{cases} S, \text{ if the switch and integrase genes are functional} \\ S', \text{ if the switch and integrase genes have been mutated} \end{cases}$$

$$G_3 = \begin{cases} P, \text{ if the synthetic protease gene is functional} \\ P', \text{ if the burdensome gene has been mutated} \end{cases}$$

$$G_4 = \begin{cases} C, \text{ if the CAT gene is functional} \\ C', \text{ if the CAT gene has been mutated or excised by the integrase} \end{cases}$$

and

$$G_5 = \begin{cases} H, \text{ if the Punisher is in a high-expression equilibrium} \\ L, \text{ if the Punisher is in a low-expression equilibrium} \\ 0, \text{ if no switch or integrase proteins are present in the cell} \end{cases}$$

Thus, for example, $x_{B'SPC:H}$ represents the number of cells in which the Punisher is in a high-expression equilibrium and only the synthetic burdensome gene has been mutated. If some statement is true for several subpopulations, we use $*$ to indicate the index entries whose values are irrelevant – e.g. referring to $j = *SPC : *$ means that a claim applies to any state population with functional switch/integrase, protease and CAT genes, regardless of the Punisher's state or of whether the synthetic constitutive burdensome gene is functional.

Importantly for our definition of $G_5$, a given Punisher equilibrium may not exist for some genetic states (e.g. most obviously, if the switch and integrase genes are non-functional, i.e. $G_2 = S'$, only a zero-expression steady state exists). However, since gene mutations and Punisher state transitions happen independently in our model, we still consider all combinations of the cell's genetic states and the Punisher's state. Namely, a cell subpopulation with an 'impossible' combination of genetic and

Punisher states can be replenished due to mutations of the cells where a given Punisher equilibrium indeed does exist. For example, the subpopulation $BS'PC : H$ is replenished when the switch gene mutates in $BSPC : H$ cells. The depletion of this $BS'PC : H$ subpopulation happens due to the concentration of the switch and integrase proteins falling, because they are degraded and diluted but no longer synthesised. This fall is equivalent to cells transitioning from the $BS'PC : H$ subpopulation to $BS'PC : L$ and then to $BS'PC : 0$.

#### S4.2 Population model ODE

The dynamics of the population state vector $\mathbf{x}$ are given by Equation (S97), which in this supplementary note is explained term-by-term.

$$\dot{\boldsymbol{x}} = (\boldsymbol{D} + \boldsymbol{A} + \boldsymbol{T} - L(\boldsymbol{x}, \boldsymbol{d}))\boldsymbol{x} \tag{S97}$$

Here, $\boldsymbol{d}$ is a vector of division rates for all cell subpopulations, ordered in the same way as in the cell count vector $\boldsymbol{x}$ (i.e. $\boldsymbol{d}_j$ is the rate of cell division in the subpopulation $j$ whose cell counts are given by the entry $\boldsymbol{x}_j$).

$\boldsymbol{D}$ is the 'cell division matrix', in which the element $\boldsymbol{D}_{j,l}$ (entry in row $j$, column $l$) is the rate of the number of cells in the subpopulation $j$ changing due to the division of mother cells in the subpopulation $l$. Upon each cell division, the mother cell ceases to exist, and two new daughter cells appear in its place. Thus, since the division rate of the cells in subpopulation $l$ is $\boldsymbol{d}_l$, division removes mother cells from the subpopulation $l$ at a rate of $\boldsymbol{d}_l$ per cell. At the same time, two new daughter cells appear at the rate $\boldsymbol{d}_l\boldsymbol{x}_l$. Overall, one cell division removes one mother cell and produces two new daughter cells, so

$$\sum_j \boldsymbol{D}_{j,l} = -\boldsymbol{d}_l + 2\boldsymbol{d}_l = \boldsymbol{d}_l \ \forall l \tag{S98}$$

The majority of daughter cells are identical to their mother, so the sum in Equation (S98) is predominated by $\boldsymbol{D}_{l,l}$. However, some divisions are accompanied by synthetic gene mutations, which makes them contribute to the subpopulation of cells whose genetic state differs from that of the mother cell. We assume that all four synthetic genes mutate independently with an equal probability $\mu$ with every cell division, and that genetic mutations do not alter the state of the

Punisher circuit. The division rate matrix's elements should thus be found recursively to account for the possibility of several genes mutating at the same time. For example, the column of $\boldsymbol{D}$ corresponding to the state $B'SPC:H$ can be retrieved according to Algorithm S1.

**Algorithm S1** Finding the division matrix column $\boldsymbol{D}_{\cdot,l}$ for the state $l = B'SPC:H$ with three functional synthetic genes.

Set $\boldsymbol{D}_{B'S'P'C':H,B'SPC:H} = \mu^3 \cdot 2\boldsymbol{d}_{B'SPC:H}$ (new cells arising with all three genes mutated)
**for** $j \in \{B'SP'C':H,\ B'S'PC':H,\ B'S'P'C:H\}$ **do**
  Set $\boldsymbol{D}_{j,B'SPC:H} = \mu^2 \cdot 2\boldsymbol{d}_{B'SPC:H} - \boldsymbol{D}_{B'S'P'C':H,B'SPC:H}$ (new cells arising with two genes mutated, but excluding the case where all three genes mutate)
**end for**
**for** $j \in \{B'SPC':H,\ B'S'PC:H,\ B'SP'C:H\}$ **do**
  Set $\boldsymbol{D}_{j,B'SPC:H} = \mu \cdot 2\boldsymbol{d}_{B'SPC:H} - \sum_{u\in\{B'SP'C':H,\ B'S'PC':H,\ B'S'P'C:H\}} \boldsymbol{D}_{u,B'SPC:H} - \boldsymbol{D}_{B'S'P'C':H,B'SPC:H}$ (new cells arising with one gene mutated, but excluding the cases where two or three genes mutate)
**end for**
Set $\boldsymbol{D}_{B'S'P'C':H,B'SPC:H} = -\boldsymbol{d}_{B'SPC:H} + 2\boldsymbol{d}_{B'SPC:H} - \sum_{u\in\{B'SPC':H,\ B'S'PC:H,\ B'SP'C:H\}} \boldsymbol{D}_{u,B'SPC:H} - - \sum_{u\in\{B'SP'C':H,\ B'S'PC':H,\ B'S'P'C:H\}} \boldsymbol{D}_{u,B'SPC:H} - \boldsymbol{D}_{B'S'P'C':H,B'SPC:H}$ (mother cell dividing, two new cells arising, but excluding the cases where one, two or three genes mutate)
**for** any other $j$ **do**
  Set $\boldsymbol{D}_{j,B'SPC:H} = 0$ (new cells like this cannot arise from state $B'SPC:H$ cells dividing)
**end for**

Besides mutating, the cell can change its genetic state if the integrase acts to excise (and thus make non-functional) the antibiotic resistance gene CAT ($C$). This affects the state vector through the term $\boldsymbol{Ax}$, where $\boldsymbol{A}$ is the integrase activity matrix. The term $\boldsymbol{A}_{j,l}$ is the rate of change of subpopulation $j$'s size as a result of integrase activity in the cells from subpopulation $l$. Let us say that the integrase activity in type $l$ cells is given by $\iota_l$, which depends on the Punisher's state, as different equilibria of the Punisher circuit stand for different integrase concentrations. Then, the column of $\boldsymbol{A}$ for the state $l$ can be found according to Algorithm S2.

The rate of cells transtitioning between different states of the Punisher equals $\boldsymbol{Tx}$, where $\boldsymbol{T}$ is the stochastic transition rate matrix, whose term $\boldsymbol{T}_{j,l}$ gives the rate of the Punisher switching from the state $l$ to the state $j$. When such a transition is possible and relevant to the overall population dynamics, its rate is determined by stochastic single-cell model simulations as per Supplementary Note S4.3.2. The element $\boldsymbol{T}_{l,l}$ captures the rate of subpopulation $l$ cell counts decreasing due to stochastic transitions, and is thus equal to minus the sum of all non-zero transition rates out of state $l$. Otherwise, we set $\boldsymbol{T}_{j,l} = 0$.

**Algorithm S2** Finding the integrase action rate matrix column $\boldsymbol{A}_{\cdot,l}$ for the generic state $l$.

```
if l = ∗∗∗∗ : 0 then
    Set A_{j,l} = 0 ∀j (no integrase present in the cell)
else if l = ∗∗∗C′ : ∗ then
    Set A_{j,l} = 0 ∀j (CAT gene already non-functional)
else
    Find l′ identical to l, but with C′ in place of C
    Set A_{l′,l} = ι_l (a cell with non-functional CAT gene is created)
    Set A_{l,l} = −ι_l (one fewer cell now has a functional CAT gene)
    for j ∉ {l′, l} do
        Set A_{j,l} = 0 (integrase action cannot create such cells)
    end for
end if
```

The final term in Equation (S97) is $-\boldsymbol{x}L$, where $L(\boldsymbol{x},\boldsymbol{d})$ is the scalar rate of all cells being diluted in the bioreactor, calculated from the subpopulations' cell counts and growth rates. We assume our reactor to be a turbidostat that regulates the cell dilution rate to maintain a constant cell density in the culture (and thus a constant overall number of cells in the bioreactor) [2]. Hence, the dilution rate is given by the overall rate of new cells appearing as a result of cell division. Since different cell subpopulations grow at different rates, $L$ is thus equal to the average of all subpopulations' rates of cell division weighted by the numbers of cells in each subpopulation. This quantity is given by Equation (S99), where $sum(\boldsymbol{x})$ is the sum of all elements in the vector $\boldsymbol{x}$.

$$L(\boldsymbol{x},\boldsymbol{d}) = \frac{\boldsymbol{x}\cdot\boldsymbol{d}}{sum(\boldsymbol{x})} \tag{S99}$$

#### S4.3 Population model parameterisation

In this supplementary note, we outline how simulations of our coarse-grained resource-aware single-cell model (described in Supplementary note S1.1) are used to determine the parameters of the population model defined in the previous two sections.

##### S4.3.1 Cell division rates, integrase activities, synthetic protein abundances

The population model in Supplementary Notes S4.1–S1.5 relies on the knowledge of cell division rates $\{\boldsymbol{d}_j\}$ and integrase activities $\{\iota_j\}$ for all cell subpopulations. In addition to this, in the main text we evaluate the productivity and durability of engineered cell populations. This is done by calculating the rate of burdensome synthetic protein production per cell $\Theta$, which requires knowing the concentration of the burdensome protein per cell $\{p_b(j)\}$ for all subpopulations.

Each of our model's subpopulations represents cells with a given set of functioning synthetic genes whose Punisher circuit is in a given steady state. Thus, for a subpopulation we can retrieve the corresponding quantities of interest by running a deterministic simulation of our single-cell resource-aware model. We find the cell's steady states by integrating model ODEs over 50 hours of simulated time – if a zero- or low-expression equilibrium for the Punisher is desired, we set the initial switch protein level to be $m_s(t = 0\ h) = 0\ nM$; to retrieve a high-expression steady state, the initial condition $m_s(t = 0\ h) = 3,000\ nM$ was used instead. The resultant steady-state growth rates, integrase concentrations and synthetic burdensome protein levels are provided in Supplementary Table S7. In the genetic states already without the CAT gene (i.e. $***C' : *$), the Punisher's state did not matter, so the deterministic simulation was run just once with the initial condition $p_s(t = 0\ h) = 0\ nM$.

Importantly for our filling of Supplementary Table S7, the population model includes some 'impossible' cell states – that is, Punisher equilibria that are in fact unachievable for a given genetic state of the cell, but still are assumed to exist in our model. They arise when a 'progenitor' cell with the same Punisher state (but a genetic state which makes this Punisher equilibirum possible) has one of its genes mutated. Naturally, deterministic simulations will not allow to retrieve non-existent steady states. Therefore, in this case, the integrase level $p_i(j)$ in such an impossible state $j$ is assumed to be equal to that in the corresponding progenitor cell. Conversely, since the synthetic protein levels and growth rates primarily depend on the burden experienced by the host cell (and the Punisher's weakly expressed genes do not make a significant contribution to it), the values of $p_b(j)$ and $\lambda_j$ are taken from the cell that is in the same genetic state as $j$ but has an actually plausible Punisher state.

Based on the cell growth rates $\{\lambda_j\}$ and integrase concentrations $\{p_i(j)\}$ from Supplementary Table S7, we can calculate the cell division rates $\{\boldsymbol{\delta}_j$ and the rates of CAT gene excision by the integrase $\{\iota_j\}$. Namely, in order to find the rate of cell division (which occurs when the cell doubles its size and protein content), the cell growth rate must be scaled by $ln(2)$ [12], which yields us:

$$\boldsymbol{d}_j = \frac{\lambda_j}{ln(2)} \quad \forall j \tag{S100}$$

Meanwhile, for the excision of the CAT gene by the integrase we recall that a functional CAT gene (with a concentration $c_{cat}$) first undergoes an integrase-mediated reversible strand exchange form an attL-attR site complex (with a concentration $c_{LRi}$), which then is irreversibly broken down either by DNA conformation changes or by plasmid replication at the rate of cell growth. This is represented by Supplementary Note S1.2's ODEs (S20)–(S21):

$$\dot{c}_{cat} = -k_{sx}^{+}\frac{p_i^4}{K_{bI}^4 + p_i^4}c_{cat} + k_{sx}^{-}c_{LRi}$$
$$\dot{c}_{LRi} = k_{sx}^{+}\frac{p_i^4}{K_{bI}^4 + p_i^4}c_{cat} - k_{sx}^{-}c_{LRi} - (k^{conf} + \lambda)c_{LRi}$$

To further simplify the system, we look at its parameters in Supplementary Table S3 and observe that the rates $k_{sx}^{+}$ and $k_{sx}^{-}$ of forward and reverse integrase-mediated strain exchange are greater than both the conformation change rate $k^{conf}$ and the typical cell growth rate $\lambda$, which in Supplementary Table S7 never exceeds 1.6. This allows us to treat the excision of the CAT gene as a one-step reaction as shown in Equation (S101)

$$\dot{c}_{cat} = -c_{cat} \cdot \frac{p_i^4}{K_{bI}^4 + p_i^4} \cdot \frac{k_{sx}^{+}}{k_{sx}^{-}} \cdot (k^{conf} + \lambda) \tag{S101}$$

Remembering that the CAT gene's copy number in our population model is not a deterministic variable but a discrete quantity, starting at 1 copy per cell (see Supplementary table S3), we can find the propensity for the excision of this one copy by the integrase in a state $j$ cell as:

$$\iota_j = \frac{p_i^4}{K_{bI}^4 + p_i^4} \cdot \frac{k_{sx}^{+}}{k_{sx}^{-}} \cdot (k^{conf} + \lambda) \tag{S102}$$

Supplementary Table S7: Growth rate and integrase and burdensome protein concentrations retrieved for different genetic states. For 'possible' cell states, the comments in brackets tell whether the quantities of interest were found by deterministic simulation. For 'impossible' genetic and Punisher state combinations, we indicate the 'possible' state combination from whose simulations these values were taken.

| Genetic state | Punisher state | Growth rate $\lambda$, $h^{-1}$ | Burdensome protein conc. $p_b$, $nM$ | Integrase conc. $p_i$, $nM$ |
|---|---|---|---|---|
| $BSPC$ | $H$ | 1.33 (simulated) | $2.03 \cdot 10^5$ (simulated) | 87.8 (simulated) |
| | $L$ | 1.34 (simulated) | $2.03 \cdot 10^5$ (simulated) | 3.52 (simulated) |
| | 0 | Irrelevant$^\sharp$ | Irrelevant$^\sharp$ | Irrelevant$^\sharp$ |
| $B'SPC$ | $H$ | 1.53 (simulated) | 0 (simulated) | 104 (simulated) |
| | $L$ | 1.53 ($B'SPC:H$) | 0 ($B'SPC:H$) | 3.52 ($BSPC:L$) |
| | 0 | Irrelevant$^\sharp$ | Irrelevant$^\sharp$ | Irrelevant$^\sharp$ |
| $BS'PC$ | $H$ | 1.33 ($BS'PC:0$) | $2.03 \cdot 10^5$ ($BS'PC:0$) | 87.8 ($BSPC:H$) |
| | $L$ | 1.33 ($BS'PC:0$) | $2.03 \cdot 10^5$ ($BS'PC:0$) | 3.52 ($BSPC:L$) |
| | 0 | 1.33 (simulated) | $2.03 \cdot 10^5$ (simulated) | 0 (simulated) |
| $B'S'PC$ | $H$ | 1.53 ($B'S'PC:0$) | 0 ($B'S'PC:0$) | 104 ($B'SPC:H$) |
| | $L$ | 1.53 ($B'S'PC:0$) | 0 ($B'S'PC:0$) | 3.52 ($B'SPC:L^\aleph$) |
| | 0 | 1.53 (simulated) | 0 (simulated) | 0 (simulated) |
| $BSP'C$ | $H$ | 1.33 (simulated) | $2.02 \cdot 10^5$ (simulated) | 101 (simulated) |
| | $L$ | 1.33 ($BSP'C:H$) | $2.02 \cdot 10^5$ ($BSP'C:H$) | 3.52 ($BSPC:L$) |
| | 0 | Irrelevant$^\sharp$ | Irrelevant$^\sharp$ | Irrelevant$^\sharp$ |
| $B'SP'C$ | $H$ | 1.52 (simulated) | 0 (simulated) | 115 (simulated) |
| | $L$ | 1.52 ($B'SP'C:H$) | 0 ($B'SP'C:H$) | 3.52 ($B'SPC:L^\aleph$) |
| | 0 | Irrelevant$^\sharp$ | Irrelevant$^\sharp$ | Irrelevant$^\sharp$ |
| $BS'P'C$ | $H$ | 1.34 ($BS'P'C:0$) | $2.03 \cdot 10^5$ ($BS'P'C:0$) | 101 ($BSP'C:H$) |
| | $L$ | 1.34 ($BS'P'C:0$) | $2.03 \cdot 10^5$ ($BS'P'C:0$) | 3.52 ($BSP'C:L^\aleph$) |
| | 0 | 1.34 (simulated) | $2.03 \cdot 10^5$ (simulated) | 0 (simulated) |

| Genetic state | Punisher state | Growth rate $\lambda$, $h^{-1}$ | Burdensome protein conc. $p_b$, $nM$ | Integrase conc. $p_i$, $nM$ |
|---|---|---|---|---|
| $B'S'P'C$ | $H$ | 1.53 ($B'S'P'C:0$) | 0 ($B'S'P'C:0$) | 115 ($B'SP'C:H$) |
| | $L$ | 1.53 ($B'S'P'C:0$) | 0 ($B'S'P'C:0$) | 3.52 ($B'SP'C:L^{\aleph}$) |
| | 0 | 1.53 (simulated) | 0 (simulated) | 0 (simulated) |
| $BSPC'$ | * | 0.34 (simulated) | $7.28 \cdot 10^4$ (simulated) | Irrelevant[§] |
| $B'SPC'$ | * | 0.35 (simulated) | 0 (simulated) | Irrelevant[§] |
| $BS'PC'$ | * | 0.34 (simulated) | $7.28 \cdot 10^4$ (simulated) | Irrelevant[§] |
| $B'S'PC'$ | * | 0.35 (simulated) | 0 (simulated) | Irrelevant[§] |
| $BSP'C'$ | * | 0.34 (simulated) | $7.27 \cdot 10^4$ (simulated) | Irrelevant[§] |
| $B'SP'C'$ | * | 0.35 (simulated) | 0 (simulated) | Irrelevant[§] |
| $BS'P'C'$ | * | 0.34 (simulated) | $7.28 \cdot 10^4$ (simulated) | Irrelevant[§] |
| $B'S'P'C'$ | * | 0.35 (simulated) | 0 (simulated) | Irrelevant[§] |

[♯]State not reachable – see the definition of the Punisher state transition rate matrix $\boldsymbol{T}$ in Supplementary Note S4.3.2.

[ℵ]Here, the progenitor state is also impossible, so we use the values for the progenitor's progenitor, i.e. the genetic state $BSPC$ and a relevant state of the Punisher.

[§]CAT gene already non-functional, so integrase activity does not matter.

##### S4.3.2 Punisher state transition rates

Besides the rates of cell division and CAT gene excision by the integrase, the population model also includes the rates of transition between the Punisher's states, which are captured by the transition rate matrix $\boldsymbol{T}$. In this matrix, the element $\boldsymbol{T}_{j,l}$ represents the rate of cells transitioning from the subpopulation $l$ to the subpopulation $j$ due to the Punisher circuit switching its state.

For most pairs of subpopulations $j$ and $l$, such transitions are impossible. For others, state transitions can also just be irrelevant: once the CAT gene has been excised, the state of the Punisher – i.e. the concentration of its integrase and switch proteins – no longer has any bearing on the cell's genetic state. These cases, for which we set $\boldsymbol{T}_{j,l} = 0$, are as follows:

1. $j$ and $l$ stand for genetically different subpopulations, since the Punisher's switching does not itself change the cell's genetic state.

2. $j = **** : 0$ and $l = **** : H$ or $j = **** : H$ and $l = **** : 0$, since the Punisher cannot 'jump' over the low-expression equilibrium which lies between the zero- and high-expression steady states

3. $j = *S' ** : L$ and $l = *S' ** : 0$ or $j = *S' ** : H$ and $l = *S' ** : L$, since the switch protein's concentration cannot increase if the switch gene is non-functional

4. $j = *S ** : 0$, since if the switch gene is functional, there is no equilibrium at the zero expression point, so no transitions to this state occur

5. $l = *S ** : 0$, since if the original state is unreachable (see Case 4 above), transition rates from it do not need to be calculated either

6. $l = ***C' : *$. If the CAT gene is already non-functional, we do not care about the Punisher's state anymore, even if it can change. Indeed, the Punisher's state primarily determines the corresponding propensity of the integrase to cut out the essential gene. Meanwhile, as discussed in Supplementary Note S4.3.1, the effect of the Punisher's state on cell growth rates is negligible.

The rates of all possible and relevant Punisher state transitions $\{\boldsymbol{T}_{j,l}\}$, were determined by stochastic simulation. This allowed to not only capture the 'true positive' switchings of the Punisher's state as it reacts to changes in gene expression burden, but also 'false positive' transitions simply occurring due to the stochasticity of gene expression. The state transition times obtained from stochastically simulated trajectories constitute a sample from the underlying probability distribution of first-passage time from one state to another. We thus assumed a Poisson distribution of such first-passage times and estimated its mean using the the maximum likelihood estimation method described in [16].

Specifically, the system was simulated deterministically to make it reach the original steady state $l$ using the initial conditions stated in Supplementary Note S4.3.1. Then, a hybrid tau-leaping simulation was run to observed the system's stochastic behaviour over $t_{j,l}^{stochsim}$ simulated hours. If during this time the integrase's concentration came within 10% of the value corresponding to the destination steady state $j$, the first time when this happened ($\tilde{t}_{j,l}^{switch}$) was recorded as the Punisher's 'switching time' for this trajectory. Such stochastic simulations were performed $N_{j,l}$ times.

Then, the mean time $\hat{t}_{j,l}^{switch}$ of switching from $l$ to $j$ was estimated from these stochastically sampled trajectories according to [16], where the Poisson distribution of the Punisher's switching times is approximated as normal. For the majority of transitions, this was calculated using Equation (S103) with $U_{j,l}$ defined as the the number of samples for which the switching did occur and a $\tilde{t}^{switch}$ value was recorded.

$$\hat{t}_{j,l}^{switch} = \frac{sum\ of\ all\ \tilde{t}_{j,l}^{switch}\ values\ recorded}{U_{j,l}} + \frac{(N_{j,l} - U_{j,l})t_{j,l}^{stochsim}}{N_{j,l}} \tag{S103}$$

In order to check the reliability of our estimates, a 95% confidence interval for our $\hat{t}_{j,l}^{switch}$ estimates was evaluated using the statistic of the switching time $\zeta(t_{j,l}^{switch})$, which is found according to Equation (S104)) and has a unit normal distribution [16]. The resultant widths of confidence intervals for the switching times, shown in Supplementary Table S9, never constituted more than 3% of the maximum likelhiood estimate, indicating that our chosen sample size $N$ and the stochastic simulation time $t^{stochsim}$ were sufficiently high to enable accurate estimation of switching times.

$$\zeta(t_{j,l}^{switch}) = \frac{w_0\sqrt{N_{j,l}(1-Q)}}{\sqrt{1 - 2w_0w_1 + Q(w_0)^2}} \tag{S104}$$

where

$$Q = exp(-t_{j,l}^{stochsim}/t_{j,l}^{switch}), \quad w_0 = \frac{\hat{t}_{j,l}^{switch} - t_{j,l}^{switch}}{t_{j,l}^{switch}}, \quad w_1 = -\frac{t_{j,l}^{stochsim}}{t_{j,l}^{switch}} \cdot \frac{Q}{1-Q} \tag{S105}$$

An exception to this scheme was made for transitions from $BSPC : H$ to $BSPC : L$ and from

$B'SPC : H$ to $B'SPC : L$, where switchings of the Punisher were recorded very rarely, i.e. the probability $Q_{j,l}$ of the Punisher remaining unswitched was much smaller than $^1/_2$. In this case, an accurate estimate for the switching rates could be calculated based solely on $U_{j,l}$, the number of samples where the switching did occur [16]. This estimate was found according to Equation (S106).

$$\hat{t}_{j,l}^{switch} = -\frac{t_{j,l}^{stochsim}}{ln(1 - {}^{U_{j,l}}/_{N_{j,l}})} \tag{S106}$$

The resultant estimated mean switching times $\{\hat{t}_{j,l}^{switch}\}$ were recorded in Supplementary Table S9, together with the specifications of our stochastic sampling schemes. The transition rate from state $l$ to state $j$ was found as the reciprocal of the estimated switching time according to Equation (S108).

$$\boldsymbol{T}_{j,l} = \frac{1}{\hat{t}_{j,l}^{switch}} \tag{S107}$$

Finally, since the Punisher's switching involves a cell transitioning to a new state **from** the old one, the element $\boldsymbol{T}_{l,l}$ represents the net transitions of cells **out** of the subpopulation $l$. Thus, it is given by Equation (S108).

$$\boldsymbol{T}_{l,l} = -\sum_{\forall j \neq l} \boldsymbol{T}_{j,l} \tag{S108}$$

Supplementary Table S9: Estimation of times taken by the Punisher to stochastically switch between its equilibria for different genetic states of the host cell. Only possible and relevant transitions are considered. In all cases, $N_{j,l} = 30,000$ sample trajectories were used.

| Genetic state | Punisher state transition | Estimated switching time $\hat{t}_{j,l}^{switch}$, $h$ | Simulation time $t_{j,l}^{stochsim}$, $h$ | 95% confidence interval width, $h$ |
|---|---|---|---|---|
| $BSPC$ | $L$ to $H$ | 10.51 | 10 | 0.30 |
| | $H$ to $L$ | 1549.40 | 10 | Not applicable$^{§}$ |
| $B'SPC$ | $L$ to $H$ | 5.31 | 10 | 0.13 |
| | $H$ to $L$ | 37495.00 | 4.9 | Not applicable$^{§}$ |
| $BS'PC$ | $H$ to $L$ | 2.59 | 10 | 0.06 |
| | $L$ to 0 | 1.49 | $4.9^{\aleph}$ | 0.03 |
| $B'S'PC$ | $H$ to $L$ | 2.40 | 10 | 0.05 |
| | $L$ to 0 | 1.27 | $6.9^{\aleph}$ | 0.03 |
| $BSP'C$ | $L$ to $H$ | 3.04 | 10 | 0.07 |
| | $H$ to $L$ | $\infty^{\daleth}$ | 10 | Not applicable$^{\daleth}$ |
| $B'SP'C$ | $L$ to $H$ | 2.63 | 10 | 0.06 |
| | $H$ to $L$ | $\infty^{\daleth}$ | 10 | Not applicable$^{\daleth}$ |
| $BS'P'C$ | $H$ to $L$ | 2.71 | 10 | 0.06 |
| | $L$ to 0 | 1.49 | $4.85^{\aleph}$ | 0.03 |
| $B'S'P'C$ | $H$ to $L$ | 2.47 | 10 | 0.06 |
| | $L$ to 0 | 1.27 | $5.62^{\aleph}$ | 0.03 |

$^{§}$Switching time estimated according to Equation (S106), so the confidence interval calculation was not applicable.

$^{\aleph}$For the sake of efficiency, instead of taking new samples, the trajectories that reached the $L$ state starting at the $H$ state were left to run until they reached the 0 state. In order to find the switching time for the $L$ to 0 transition, these trajectories were truncated so as to start at the point where the Punisher was in the $L$ state, hence $t_{j,l}^{stochsim} < 10\ h$.

$^{\daleth}$Among $30,000$ trajectories, not a single one experienced a state transition during the simulation. Transitions were therefore assumed to never occur.
